# Dynamics of the 4D genome during lineage specification, differentiation and maturation *in vivo*

**DOI:** 10.1101/763763

**Authors:** A. Marieke Oudelaar, Robert A. Beagrie, Matthew Gosden, Sara de Ornellas, Emily Georgiades, Jon Kerry, Daniel Hidalgo, Joana Carrelha, Arun Shivalingam, Afaf H. El-Sagheer, Jelena M. Telenius, Tom Brown, Veronica J. Buckle, Merav Socolovsky, Douglas R. Higgs, Jim R. Hughes

## Abstract

Mammalian gene expression patterns are controlled by regulatory elements, which interact within Topologically Associating Domains (TADs). The relationship between activation of regulatory elements, formation of structural chromatin interactions and gene expression during development is unclear. We developed Tiled-C, a low-input Chromosome Conformation Capture (3C) approach, to study chromatin architecture at high spatial and temporal resolution through *in vivo* mouse erythroid differentiation. Integrated analysis of matched chromatin accessibility and single-cell expression data shows that regulatory elements gradually become accessible within pre-existing TADs during early differentiation. This is followed by structural re-organization within the TAD and formation of specific contacts between enhancers and promoters. In contrast to previous reports, our high-resolution data show that these enhancer-promoter interactions are not established prior to gene expression, but formed gradually during differentiation, concomitant with progressive upregulation of gene activity. Together, these results provide new insight into the close, interdependent relationship between chromatin architecture and gene regulation during development.

## Introduction

Enhancers are non-coding regulatory elements required for precise control of gene expression during mammalian development. The interaction of active enhancer elements with their target genes is constrained by Topologically Associating Domains (TADs), ~0.1-1 Mb self-interacting regions that are usually delimited by convergent binding sites for the zinc-finger protein CTCF, so-called boundary elements^1–3^. However, the relationship between enhancer activation, formation of intra-TAD chromatin interactions and gene activation are not completely understood^4–6^. To better understand how genome structure relates to function, it is important to characterize the three-dimensional nuclear architecture of the genome at higher resolution, and to determine how it changes during differentiation and development.

It has been shown that TAD boundaries are generally established early in development and remain relatively stable during differentiation^7–10^. By contrast, interactions within TADs are extensively restructured in differentiating cells, which involves the formation of specific interactions between enhancers and promoters^11–15^. It has been suggested that this reorganization precedes gene activity and that enhancer-promoter interactions are formed prior to gene expression^16–18^. However, due to limitations in temporal and/or spatial resolution in previous studies, it is not known precisely when such interactions are formed during development. The detailed order of events and precise relationship between chromatin architecture and activation of regulatory elements and genes therefore remain unclear. For example, it is possible that enhancers and promoters form limited interactions prior to gene activation due to changes in the general self-interactivity of the TAD during differentiation, but that strong upregulation of gene expression is associated with more specific, subtle changes in conformation that will only be detected in data with sufficient resolution and sensitivity.

To better understand the relationship between chromatin architecture and gene expression, it is crucial to characterize chromatin structures at high resolution in pure, primary cell populations representing relevant developmental stages. This has been hampered by the lack of high-resolution 3C methods that are suitable for the analysis of limited numbers of primary cells. Therefore, we developed Tiled-C, a new 3C-based approach^19^, which can generate high-resolution contact matrices of selected regions of interest from as few as 2,000 cells and thereby allows for the analysis of cell populations that have previously been inaccessible.

We have used Tiled-C to study the chromatin architecture of key erythroid genes through sequential stages of *in vivo* erythroid differentiation in the mouse, including highly purified hematopoietic stem and progenitor cells. In addition, we have generated matched chromatin accessibility and single-cell expression data. We examined six loci, including the α-globin, *Slc25a37*, *Tal1*, *Cd47*, *Cpeb4* and *Btg2* genes, and focused our analyses on the α-globin genes, because the regulatory elements in this locus are extremely well-characterized. We find that the TAD encompassing the α-globin genes is already present in hematopoietic stem cells. We also find that the first steps in gene activation occur in early committed erythroid progenitors and involve opening of the α-globin enhancers, which become accessible prior to both chromatin reorganization and activation of α-globin RNA expression. Subsequent chromatin reorganization involves the appearance of smaller self-interacting domains within the larger TAD, in which specific interactions between enhancers and promoters are formed. In contrast to the current literature^16–18^, we find that these enhancer-promoter interactions do not precede upregulation of gene activity, but are formed gradually and concomitantly with progressive activation of α-globin expression. Importantly, we find a similar order of events at other erythroid gene loci. Therefore, our data demonstrate that – at this improved level of resolution – chromatin architecture and gene activation are more tightly linked than previously appreciated. Together, these findings provide new insights into the mechanisms contributing to the establishment of tissue-specific chromatin structures during development.

## Results

### Tiled-C generates high-resolution contact matrices from small numbers of cells

We developed Tiled-C, a 3C technique which generates deep, high-resolution contact matrices of genomic regions of interest. Tiled-C maximizes library complexity by employing a single-tube protocol for 3C library preparation, which minimizes losses during the procedure^20^. This is combined with enrichment derived from the efficient Capture-C technology, which allows for up to a million-fold enrichment of restriction fragments of interest^21,22^. While Capture-C targets individual restriction fragments as viewpoints, Tiled-C uses a panel of capture oligonucleotides tiled across all restriction fragments of specified genomic regions to efficiently enrich for contacts within this region. This allows for deep, targeted sequencing of chromatin interactions within regions of interest and thus for the generation of high-resolution contact matrices at unprecedented depth, across multiplexed samples and genomic regions. Tiled-C therefore combines the ability of all vs all methods such as Hi-C^23^ to map large-scale chromatin structures including TADs, and the ability of one vs all methods such as 4C^24,25^ and Capture-C^21,22^ to robustly identify enhancer-promoter interactions within TADs in detail (Supplementary Figure 1,2). To validate the Tiled-C approach, we compared Tiled-C data to the deepest currently available *in situ* Hi-C datasets (mouse ES cells^9^; Figure 1a). Tiled-C data at this region was ~28-fold higher in depth and required ~19-fold less sequencing (Supplementary Table 1). To further demonstrate the capabilities of Tiled-C, we performed experiments in which we multiplexed across several key gene loci in both primary mouse erythroid cells and ES cells, to characterize their tissue-specific configurations in depth (Figure 1b,c)

**Figure 1:**
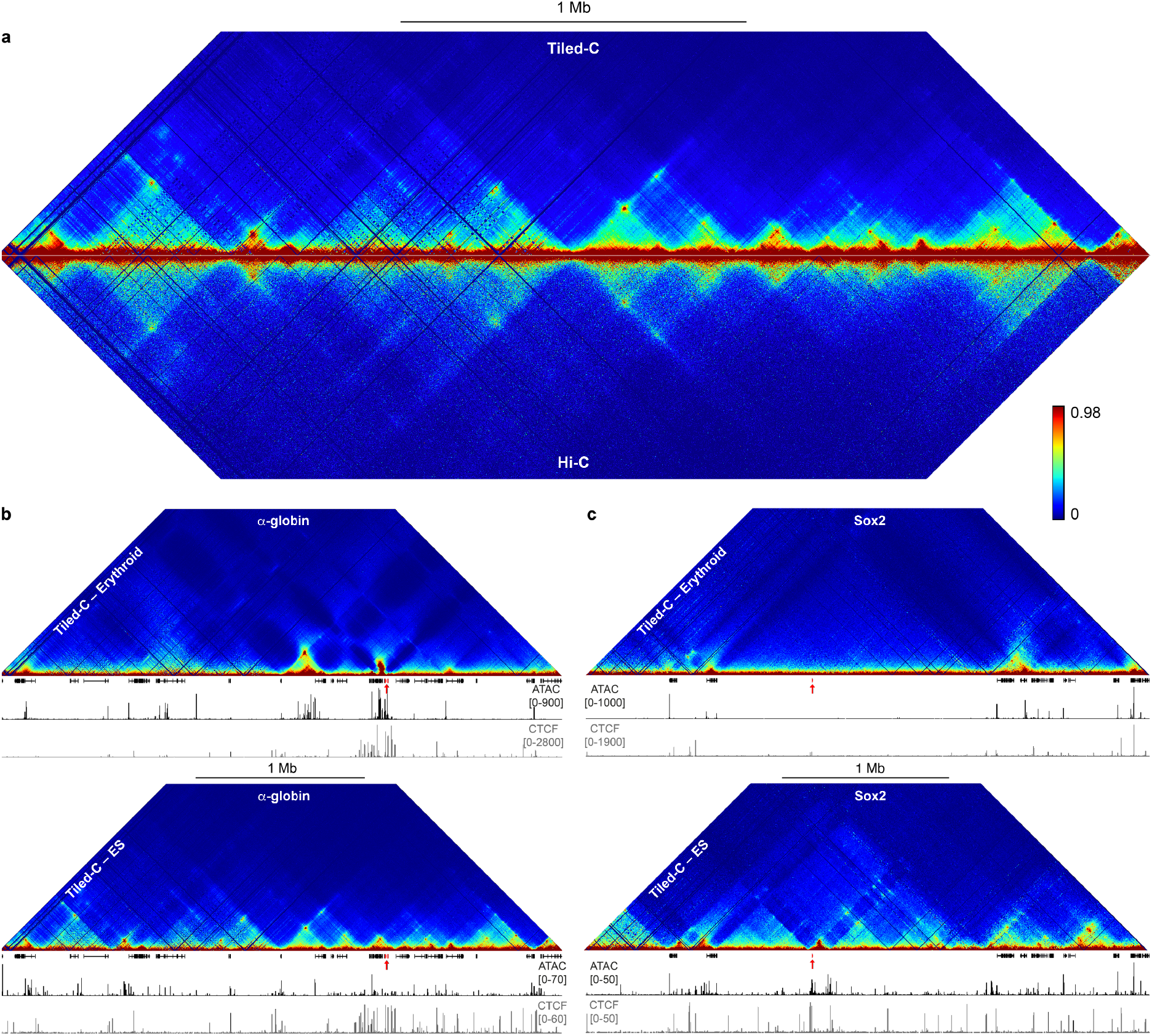
Tiled-C generates deep all vs all 3C data at regions of interest. **(a)** Comparison of Tiled-C and Hi-C contact matrices at 2 kb resolution in mouse ES cells. Contact frequencies represent normalized, unique interactions in 3 and 4 replicates for Tiled-C and Hi-C data, respectively. Coordinates (mm9): chr11:29,902,000-33,228,000. **(b)** Tiled-C contact matrices of ~3.3 Mb spanning the α-globin locus in primary mouse erythroid cells (top) and ES cells (bottom) at 2 kb resolution. Contact frequencies represent normalized, unique interactions in 3 replicates. Gene annotation (α-globin genes highlighted in red), open chromatin (ATAC) and CTCF occupancy are shown below the matrices. Coordinates (mm9): chr11:29,902,000-33,228,000. **(c)** Tiled-C contact matrices of ~3.4 Mb spanning the Sox2 locus in primary mouse erythroid cells (top) and ES cells (bottom) at 5 kb resolution. Contact frequencies represent normalized, unique interactions in 4 replicates. Gene annotation (Sox2 gene highlighted in red), open chromatin (ATAC) and CTCF occupancy are shown below the matrices. Coordinates (mm9): chr3:33,200,000-36,565,000.

Current methods such as Targeted Chromatin Capture (T2C)^26^, Capture Hi-C^27^ and HYbrid Capture Hi-C (Hi-C^2^)^28^ also use an oligonucleotide capture procedure to enrich 3C or Hi-C libraries for regions of interest (Supplementary Table 2). These methods require millions of cells per sample. In contrast, as Tiled-C is specifically designed to maximize the efficiency of the experimental procedure, its increased sensitivity allows for the generation of reproducible, high-resolution contact matrices from as few as 2,000 cells (Figure 2, Supplementary Figure 3-4, Supplementary Table 3), thereby enabling the analysis of previously intractable primary cell types. This is critical for the investigation of 4D (3D structure through developmental time) genome organization, as cell numbers become extremely limiting at early stages of development.

**Figure 2:**
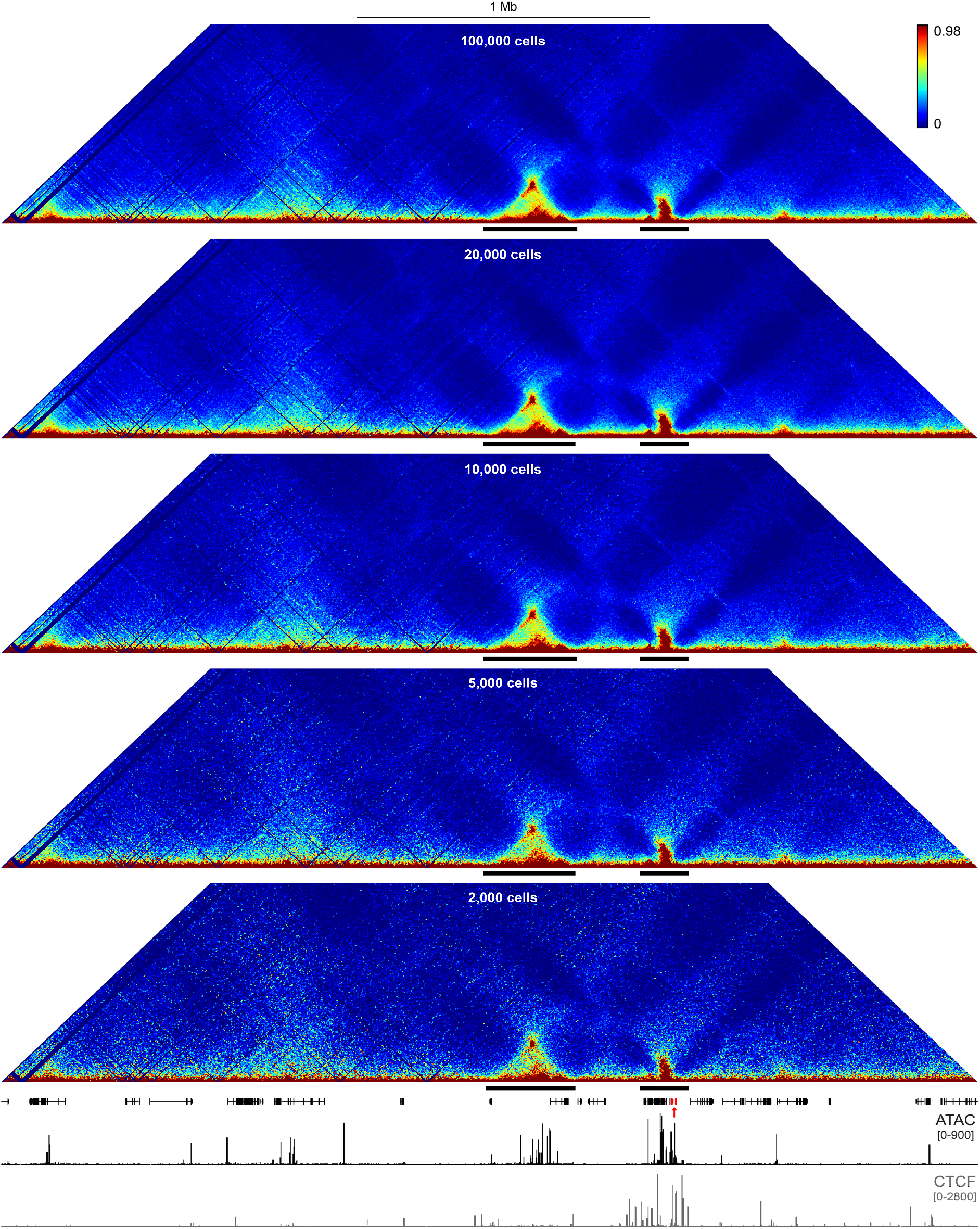
Tiled-C generates high-resolution contact matrices from small numbers of cells. Tiled-C contact matrices of ~3.3 Mb spanning the α-globin locus generated from small aliquots of primary mouse erythroid cells at 5 kb resolution. Contact frequencies represent normalized, unique interactions in 3 replicates. TADs are indicated below each matrix with a black bar. Gene annotation (α-globin genes highlighted in red), open chromatin (ATAC) and CTCF occupancy are shown below the matrices. Coordinates (mm9): chr11:29,900,000-33,230,000.

We used Tiled-C to study changes in chromatin structure associated with gene activation in primary cells during *in vivo* mouse erythropoiesis. We initially focused on the αα-globin cluster because the α-globin genes and their regulatory elements are extremely well-characterized. The mouse α-globin cluster comprises the duplicated adult α-globin genes *Hba-1* and *Hba-2*, as well as the embryonic gene *Hba-x*, and two genes of unknown function *Hbq-1* and *Hbq-2*. These genes are located in a TAD, which also contains five additional genes upstream of the α-globin cluster: *Nprl3*, *Mpg*, *Rhbdf1*, *Snrnp25* and *Il9r*. The α-globin genes are regulated by five erythroid-specific enhancer elements (R1-R4 and Rm), which classify as a super-enhancer^29^. In terminally differentiating erythroblasts these enhancers interact with the gene promoters in a sub-TAD flanked by multiple CTCF-binding elements, which are predominantly in a convergent orientation^30,31^ (Supplementary Figure 1).

### Isolation of primary cell populations from sequential stages of *in vivo* erythropoiesis

Using fluorescence-activated cell sorting (FACS), we isolated cells at sequential stages of erythroid differentiation directly from mouse fetal livers (Figure 3a,b). This allowed us to analyze highly purified primary cells through *in vivo* erythropoiesis. The S0-low cell population consists of early progenitors, predominantly Burst-Forming Unit-Erythroid (BFU-E) cells. S0-medium consists primarily of early Colony-Forming Unit-Erythroid (CFU-E) cells, while S1 and S2 contain the last CFU-E cell division before terminal differentiation^32,33^. S3 through S5 consist of terminally differentiating erythroblasts in progressively more mature states. Because erythroid cells enucleate in the final stages of differentiation, we have focused our analyses on stages S0 through to S3. In vitro, differentiation from S0 cells to S1 cells takes about 10 hours, and differentiation to S3 cells takes an additional 10 hours.

**Figure 3:**
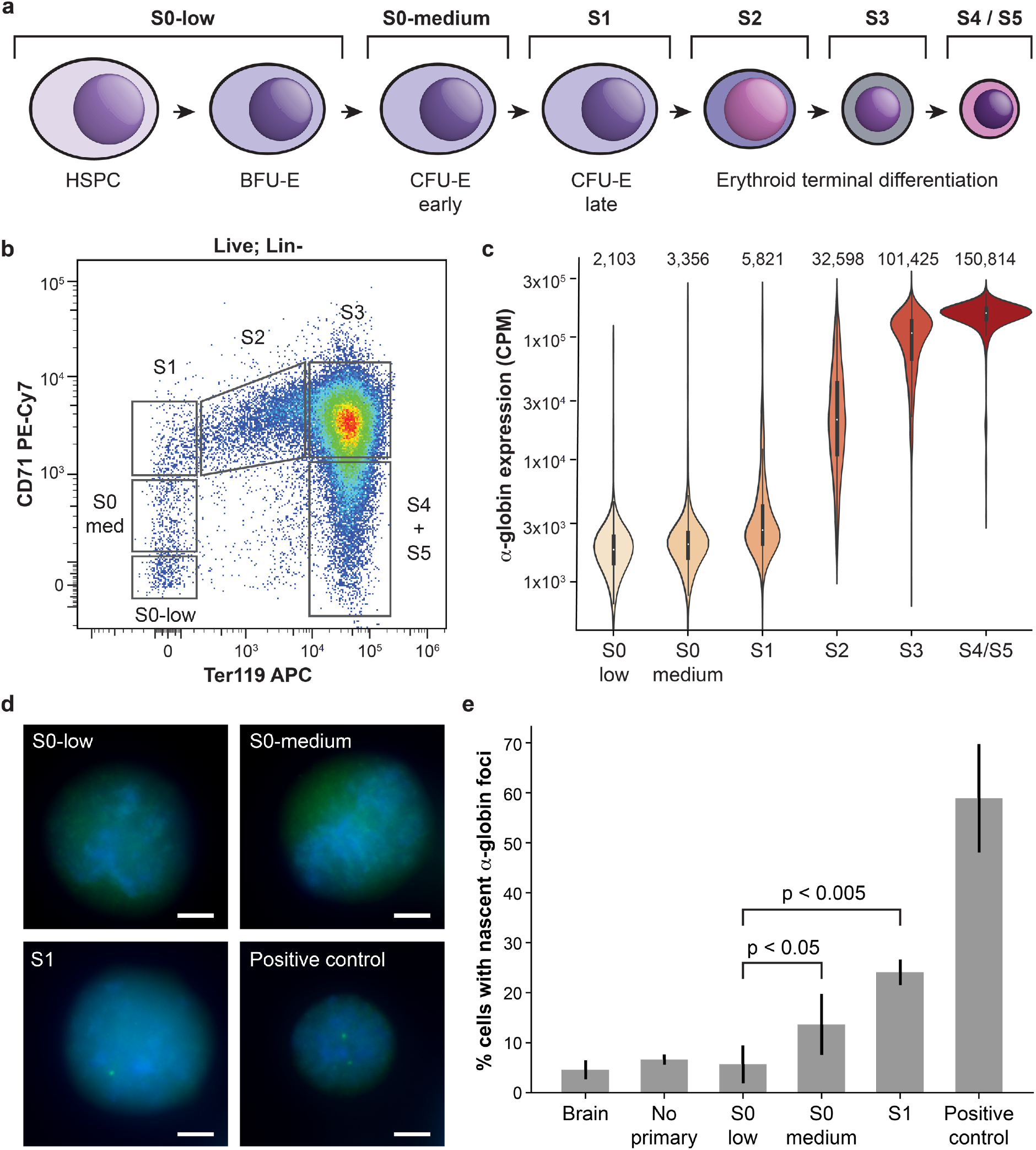
Expression of α-globin is gradually upregulated during in vivo erythroid differentiation. **(a)** Scheme of erythroid differentiation showing the various populations analyzed. **(b)** Example FACS plot showing the gating strategy used to isolate erythroid progenitors from mouse fetal liver. **(c)** Expression (in counts per million) of α-globin transcripts in each population as determined by single-cell RNA-seq. Mean expression for each population is given at the top of each bar. **(d)** Representative RNA-FISH images showing detection of nascent α-globin transcripts in sorted early erythroid progenitors. Scale bar is 3 μM for each image. **(e)** RNA-FISH quantification, showing the mean ± s.d. of 3 independent experiments (except for brain and “no primary” negative controls, which have n=2). P values were calculated by two-tailed paired T-tests.

### Characterization of gene expression through in vivo erythroid differentiation by single-cell RNA-seq

We used Cellular Indexing of Transcriptomes and Epitopes by Sequencing (CITE-seq)^34^, a variant of single-cell RNA-seq, to characterize gene expression in the isolated cell populations, generating the first dataset to include the full course of *in vivo* erythroid differentiation through to terminal differentiation in the mouse (Supplementary Figure 5). This dataset shows that α-globin is expressed at basal levels in the S0 populations. Expression of α-globin dramatically increases during the S2 stage and plateaus at S3, however the earliest cells showing elevated expression of α-globin are found in the S1 stage (Figure 3c). To validate that erythroid-specific α-globin upregulation begins at S1, we performed RNA-FISH to detect nascent transcription in FACS-sorted primary cells (Figure 3d). We detect a small increase in nascent transcription from S0-low to S0-medium cells and confirm a robust increase in expression from S0 to S1 cells (P < 0.005 by paired T-test; Figure 3e).

### Upregulation of gene expression is associated with progressive changes in chromatin architecture

We used the Assay for Transposase-Accessible Chromatin using sequencing (ATAC-seq)^35^ to profile chromatin accessibility in these stages. Interestingly, we find that both enhancer and promoter elements are accessible prior to the onset of erythroid-specific gene expression, and that the degree of accessibility gradually increases, concomitant with upregulation of α-globin expression (Figure 4, Supplementary Figure 6). Tiled-C shows that a TAD structure encompassing the α-globin locus is present at the earliest stage (S0-low), prior to the formation of weak enhancer-promoter interactions in the S0-medium stage (Figure 4, Supplementary Figures 7,8). These enhancer-promoter interactions strengthen in the subsequent S1 and S2 stages, accompanied by increases in α-globin expression and accessibility. In the S3 stage, where chromatin accessibility and expression reach their maximum levels, we observe a further increase in enhancer-promoter interactions in a sub-compartmentalized chromatin structure similar to that observed in primary erythroblasts derived from mature spleen tissue (Figure 1, Supplementary Figures 1,2). This smaller sub-compartmentalized structure, which forms within the pre-existing TAD, is delimited by convergent CTCF-binding elements that flank the α-globin enhancers and genes (Figure 4, Supplementary Figure 1). We have previously shown that these CTCF-binding elements are functionally important to restrict the interactions of the α-globin enhancers and prevent other genes within the TAD, but outside of the sub-compartmentalized structure, from being upregulated^31,36^. This suggests that this smaller erythroid-specific domain is likely formed by similar CTCF-dependent mechanisms as TADs, although it is smaller in size (~70 kb) and has very high internal interaction frequencies compared to typical TADs.

**Figure 4:**
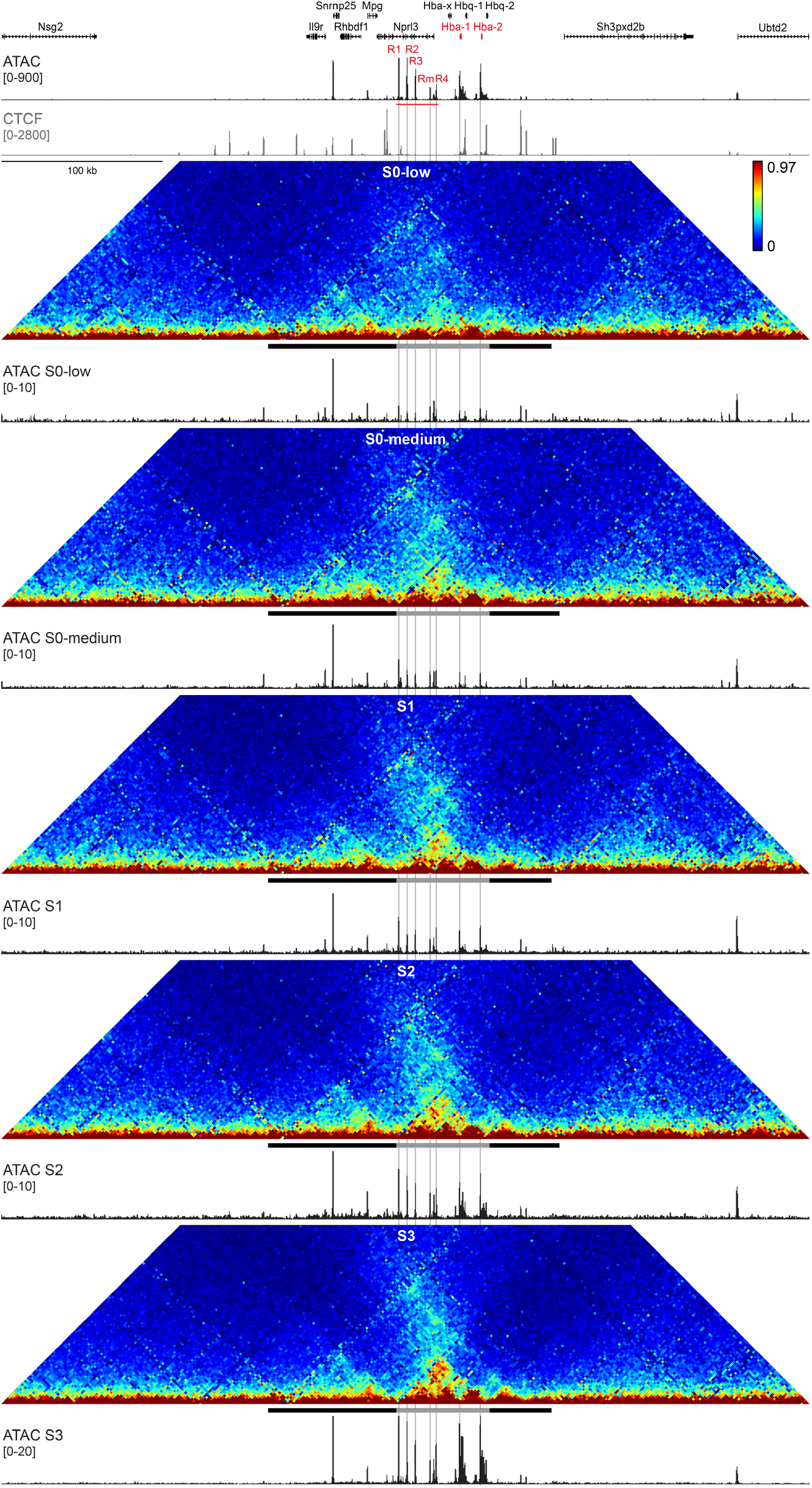
Upregulation of α-globin expression correlates with increased chromatin accessibility and enhancer-promoter interactions. Tiled-C contact matrices of 500 kb spanning the mouse α-globin locus in sequential stages of *in vivo* erythroid differentiation at 2 kb resolution. Contact frequencies represent normalized, unique interactions in 2 replicates. The black and grey bar below each matrix represent the pre-existing TAD and erythroid-specific sub-TAD, respectively. Matched open chromatin (ATAC) profiles are shown underneath the matrices and represent normalized data from 3 S0-low, S0-medium and S1 replicates and 2 S2 and S3 replicates. The ATAC profiles are shown at different scales to highlight changes in accessibility in early stages of differentiation. Gene annotation (α-globin genes highlighted in red), open chromatin (ATAC; α-globin enhancers highlighted in red) and CTCF occupancy in mature mouse erythroblast cells are shown at the top. Coordinates (mm9): chr11:31,900,000-32,400,000.

Since both accessibility and the encompassing TAD structure are present prior to erythroid-specific α-globin expression, we purified early hematopoietic progenitor populations to investigate when in differentiation these features are established. Interestingly, we find that the pre-existing TAD containing the α-globin locus is already present in hematopoietic stem cells, despite four out of five enhancers and both promoters being inaccessible at this stage (Supplementary Figure 9).

To examine whether a similar order of events operates at other gene loci, we examined the chromatin architecture of five additional erythroid gene loci through erythropoiesis: *Slc25a37*, *Tal1*, *Cd47*, *Cpeb4* and *Btg2*. In each of these loci, we find that regulatory elements are accessible prior to gene activation and gradually increase in accessibility as gene expression is upregulated (Supplementary Figures 10-12). We also find that these elements interact at basal levels in a pre-existing TAD structure prior to gene expression. However, specific interactions between the regulatory elements progressively increase during differentiation as erythroid-specific gene activity increases (Figure 5). These results confirm that specific regulatory interactions do not precede gene activation, but are formed gradually and concomitant with upregulation of gene expression.

**Figure 5:**
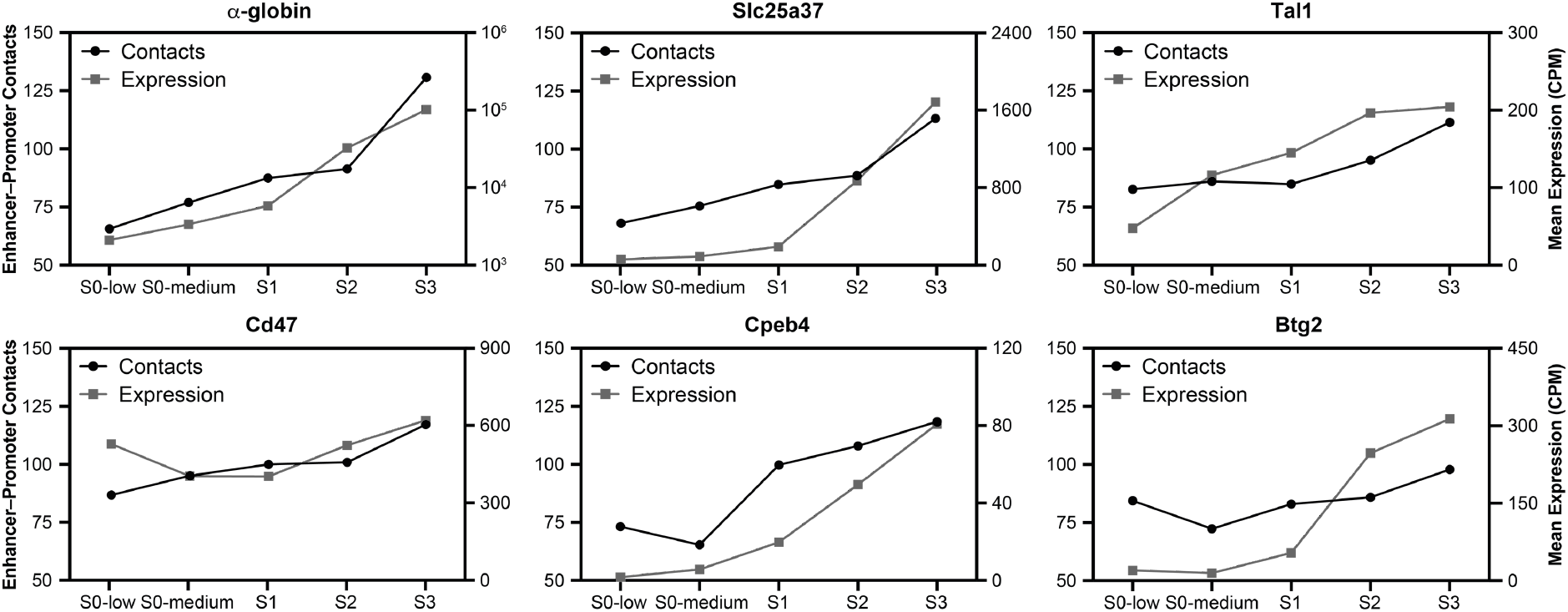
Enhancer-promoter interactions are formed progressively during erythropoiesis and correlate with upregulation of gene expression. Quantification of enhancer-promoter interactions and gene expression during in vivo erythropoiesis for six erythroid gene loci. Contact frequencies (black circles; left Y-axis) represent unique interactions normalized for the total number of contacts in the matrix. Expression counts (grey squares; right Y-axis) represent mean expression in counts per million (CPM) for each population as determined by single-cell RNA-seq. Expression of α-globin, *Slc25a37* and *Tal1* is upregulated early in differentiation, concomitant with increased enhancer-promoter interactions. *Cd47* is expressed in hematopoietic stem cells and upregulated later in erythroid differentiation when enhancer-promoter interactions are strengthened. *Cpeb4* and *Btg2* become robustly expressed in the S1 stage and are further upregulated as enhancer-promoter interactions increase later in differentiation.

## Discussion

The mouse α-globin cluster has a long history as model locus for studying gene regulation during differentiation and development^37^. Previous analysis of transcription factor binding at the α-globin locus has shown that lineage commitment and differentiation are driven by sequential appearance of key transcription factors^38^, consistent with the gradual increase in chromatin accessibility at the regulatory elements we describe here. In addition, previous 3C studies of both the α- and β-globin loci in erythroid cell lines have demonstrated that interactions between enhancers and promoters are tissue-specific^39,40^. More recently, the tissue-specific conformation of the α-globin locus has also been described based on super-resolution imaging of two stages of *ex vivo* erythroid differentiation^41^

A limitation of the current body of work on chromatin organization during differentiation and development – both at the globin clusters and other gene loci – is that experiments have been performed at low spatial and temporal resolution and predominantly *in vitro*. Previous studies have therefore not been able to identify at what point in differentiation tissue-specific interactions between enhancers and promoters are established, nor how the formation of these interactions relates to changes in gene expression. Progress in this area has been limited by a lack of techniques capable of generating high-resolution interaction data from the small numbers of cells available in developmentally relevant primary cell populations.

To overcome this hurdle, we have developed Tiled-C, a new 3C-based approach which can generate high-resolution interaction data from as few as 2,000 cells and thus enables analysis of cell populations that have previously been inaccessible. We have used Tiled-C, in combination with ATAC and single-cell RNA-seq, to study the dynamic chromatin architecture and expression of the α-globin cluster through *in vivo* erythropoiesis. Our data show that the boundaries of the TAD containing the α-globin cluster are established in hematopoietic stem cells – prior to activation of the regulatory elements and genes within the domain – and maintained during further differentiation. This is consistent with previous reports which have shown that TADs are relatively stable during differentiation and development^7–10^. In contrast to the current literature^16–18^ however, the higher resolution of our data has allowed us to show that sub-compartmentalization of the large TAD into smaller domains and the subtle structural changes that strengthen specific enhancer-promoter interactions both occur gradually during terminal erythroid differentiation, concomitant with progressive upregulation of gene activity (Figure 6). Interestingly, initial chromatin accessibility is detectable at the regulatory elements of the α-globin locus prior to conformational change and gene expression activation, although accessibility also increases gradually during differentiation. In addition to the α-globin cluster, we demonstrate the same order of events across five other erythroid gene loci.

**Figure 6:**
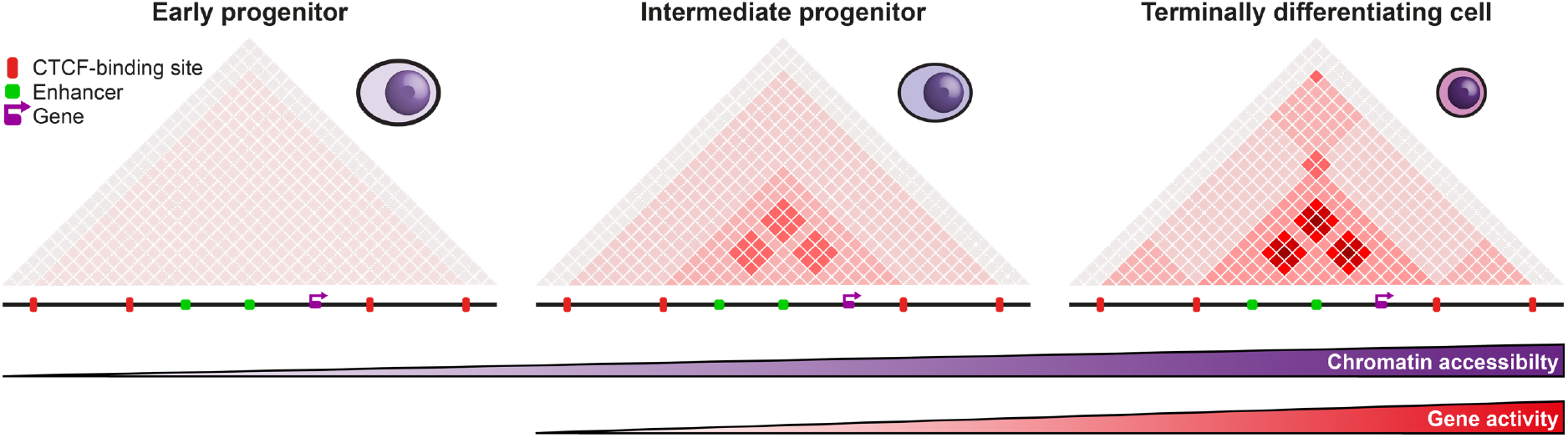
Graphical summary. Based on our findings, we propose a model in which TADs are established very early in differentiation, prior to activation of the domain. During lineage commitment, tissue-specific open chromatin sites are established within this domain. This is followed by sub-compartmentalization of the TAD into smaller domains, in which enhancers and promoters form basal interactions. Through differentiation, accessibility and specific interactions between enhancers and promoters are gradually increased, concomitant with upregulation of gene expression.

It is likely that the formation of specific sub-domains within TADs early in differentiation facilitates interactions between enhancers and promoters to prime loci for gene activation. It has recently been shown for the globin loci that gene activation is associated with the formation of higher-order hub-like structures, in which multiple enhancers and promoters form simultaneous, specific interactions^30,42^. Our data suggest that these structures may only be formed in the final stages of differentiation, when chromatin accessibility and interactions between enhancers and promoters are strongest, and may be important to achieve maximal gene expression. This is further supported by recent live imaging experiments in Drosophila, in which gene activation only occurred upon the formation of tight associations between enhancers and a gene promoter, and not after induced enhancer-promoter proximity resulting from interactions between insulator elements^43^.

This model implies that there are multiple processes contributing to the formation of specific chromatin structures associated with gene activation. A pre-existing TAD encompassing the α-globin locus is formed prior to and thus independent of activation of the regulatory elements within the domain. This is likely driven by tissue-invariant loop extrusion mediated by cohesin and constitutive CTCF-binding elements^44^. During differentiation, chromatin accessibility increases, and a smaller sub-domain is formed within this TAD. We have previously shown that deletion of the CTCF-binding sites at the base of this sub-domain causes it to expand and leads to aberrant expression of the neighboring genes^31,36^.

This indicates that sub-compartmentalization is dependent on these CTCF-binding sites and implies that its formation is mediated by loop extrusion. Since the CTCF-binding sites are constitutively occupied, erythroid-specific compartmentalization is likely driven by increased rate or processivity of loop extrusion in this region during differentiation. As we have previously observed erythroid-specific accumulation of cohesin at the α-globin enhancers^31^, it is possible that this is mediated by increased cohesin recruitment at the activated regulatory elements. This is further supported by studies showing that cohesin co-localizes with transcription factors across the genome^45,46^.

The initial appearance of chromatin accessibility at α-globin regulatory elements occurs early in differentiation and significantly precedes the onset of specific enhancer-promoter interactions. This indicates that chromatin opening can occur independently of larger scale chromatin reorganization, yet further increases in accessibility do occur alongside the establishment and progressive strengthening of enhancer-promoter interactions, suggesting only a partial decoupling. This suggests that active regulatory elements are not required for the establishments of TADs, consistent with our previous work showing that deletion of the α-globin enhancers has no impact on the formation of the α-globin TAD. These deletions do affect specific enhancer-promoter interactions^29,41^, reinforcing that regulatory elements do play a role in the formation of tissue-specific chromatin structures, possibly mediated by interactions between the multi-protein complexes bound at these elements.

In conclusion, our dissection of the chromatin architecture of a well-understood gene locus through *in vivo* erythroid differentiation demonstrates that chromatin architecture and gene activation are tightly linked during development and provides new insights into the distinct mechanisms contributing to the establishment of tissue-specific chromatin structures. Importantly, Tiled-C provides an approach that enables such detailed analysis in cell types that were previously intractable.

## Materials and Methods

### Cells

#### Mature erythroid cells

Mature primary Ter 119+ erythroblasts were obtained from spleens of female C57BL/6 mice treated with phenylhydrazine as previously described^22^.

#### Mouse ES cells

Mouse ES cells were cultured and harvested as previously described^22^.

#### Erythroid progenitors

Primary erythroid progenitor cells were isolated from fetal livers, which were freshly isolated at e12.5-e13.5 from C57BL/6 mouse embryos. 5-15 livers were pooled together for each experimental replicate, mechanically dissociated in staining buffer (PBS, 0.2% BSA, 5mM Glucose) and strained through a 30-μm strainer. Cells were immunostained at 4 °C in the presence of rabbit IgG (200 μg/ml, Jackson Laboratories 015-000-003) to block Fc receptors. To enrich for early erythroid progenitors, cells were first stained with 5 μg/ml biotin-conjugated anti-Ter119 (BD 553672) for 30 minutes, before magnetic depletion using streptavidin nanobeads (BioLegend Mojosort 480016) following the manufacturer’s instructions. Cells were then incubated with 0.5 μg/ml APC-conjugated streptavidin (BD 553672), 0.33 μg/ml PE-Cy7-conjugated anti-CD71 (BD Biosciences) and a panel of 5 FITC-conjugated lineage antibodies (anti-CD41, anti-CD45R, anti-CD3e, anti-CD11b and anti-Ly-6G/6C, all at 1 μg/ml; BD 553848, 553087, 553061, 557396 and 553126, respectively) for 45 minutes. Cells were then resuspended in FACS running buffer (staining buffer plus 2 mM EDTA). 0.66 μg/ml Hoechst was added immediately prior to sorting in order to distinguish live cells. Cells were sorted into Eppendorf tubes containing 500 μl RPMI supplemented with 10% FCS using a BD FACSAria™ Fusion machine with a 100 μM nozzle size.

#### Hematopoietic progenitors

Hematopoietic stem and progenitor cells (HSPCs), lymphoid-primed multipotential progenitors (LMPPs) and multipotent progenitors (MPPs) were stained and FACS sorted from the bone marrow of 12-week old female C57BL/6 mice as previously described^47^.

#### Replicates

The presented Tiled-C data derived from mature splenic erythroblasts and ES cells represent biological triplicates produced from separate mice or culture flasks, respectively. The presented Tiled-C data derived from hematopoietic and erythroid progenitor populations represent biological duplicates, with the exception of the S1 stage, for which we used a single biological replicate to generate technical duplicates. The presented ATAC data derived from hematopoietic and erythroid progenitor populations represent biological triplicates for the S0-low, S0-medium and S1 populations, biological duplicates for the S2 and S3 populations, and single replicates for the hematopoietic progenitor populations. The presented RNA-FISH data represent biological triplicates except for the brain and no-primary-antibody negative control, which represent biological duplicates.

#### Ethics

All protocols were approved through the Oxford University Local Ethical Review process and all experimental procedures were performed in accordance with European Union Directive 2010/63/EU and/or the UK Animals (Scientific Procedures) Act, 1986.

### Tiled-C

#### Rationale

Tiled-C is a hybrid of the all vs all 3C methods, such as Hi-C^23^, and the one vs all methods, such as 4C^24,25^ and Capture-C^21,22^. Tiled-C generates all vs all contact matrices of specified genomic regions and thus combines an unbiased all vs all view with the ability to target regions of interest, without the need to sequence chromatin interactions genome-wide. Tiled-C has similarities to 5C^48^ and Targeted Chromatin Capture (T2C)^26^. However, Tiled-C allows for reliable PCR duplicate filtering based on random sonication ends and uses more efficient capture enrichment, and is therefore able to generate data at higher resolution and depth. Tiled-C also has similarities to Capture Hi-C^27^ and HYbrid Capture Hi-C (Hi-C^2^)^28^, which use oligonucleotide capture to enrich Hi-C libraries for regions of interest (Supplementary Table 2). The main differences are that these methods enrich a biotinylated Hi-C library, while Tiled-C enriches a 3C library generated with an optimized procedure to retain maximal library complexity, which is critical for the analysis of small cell numbers. In addition, Tiled-C uses an efficient capture oligonucleotide design targeted directly to the ends of all restriction fragments present in the region of interest and an efficient capture enrichment procedure enabling up to a million-fold enrichment of restriction fragments of interest. The combination of high library complexity and efficient enrichment in Tiled-C enables high-resolution data generation at great depth and makes the method suitable for the analysis of small cell numbers. Moreover, enriching for targeted regions of interest substantially decreases sequencing costs. We should note however that the synthesis of large amounts of capture oligonucleotides can also be expensive. We therefore believe that Tiled-C is particularly useful for researchers who are interested in studying genomic regions of interest in multiple replicates and conditions and/or in primary cells with limited availability.

#### Oligonucleotide design

Tiled-C uses a panel of 70 bp oligonucleotides to enrich for regions of interest. The oligonucleotide sequences are designed complementary to both ends of each individual restriction fragment present in the region of interest. We use stringent BLAT-based filtering to ensure that the panel of oligonucleotides does not contain repetitive sequences that would decrease specific enrichment of the region of interest. To help users design capture oligonucleotides, we have developed a user-friendly python package (https://oligo.readthedocs.io/en/latest/). An overview of the oligonucleotide designs for the α-globin, Sox2 and Tal1 loci is shown in Supplementary Table 4.

#### Oligonucleotide synthesis

The enrichment step in the Tiled-C protocol can be performed with single-stranded or double-stranded biotinylated capture oligonucleotides (Supplementary Figure 4, Supplementary Table 5).

We produced panels of single-stranded oligonucleotides on an in-house Combimatrix CustomArray B3 DNA synthesiser (B3Synth_v25.1 software) using CustomArray 12K Blank Slides (CustomArray Inc., PN: 2000100-Oligo pool Application). All the probe sequences were designed to be 70 bases in length and were placed at random positions on the microarray for synthesis using Layout Designer (v4.3.1). Synthesis of oligonucleotide probe sequences occurred on individual electrodes present on the semiconductor surface of the microarray by phosphoramidite chemistry in the 3’ to 5’ direction using standard software oligonucleotide pool synthesis settings and reagents prepared according to the manufacturer’s protocols. Each sequence was synthesized in triplicate. After the synthesis of the unmodified oligonucleotide, 5’-biotin was added using a double coupling cycle with an extended 15 minutes coupling time. The final detritylation step was performed manually using the software by incubating the slides with TCA deblock (4 × 30s incubations) before washing the slide with acetonitrile four times and drying under argon. Oligonucleotides were then cleaved and deprotected on a stripping clamp system provided by the manufacturer using concentrated aqueous ammonia at 65 °C for 18 h. After cooling, the solution was recovered and the ammonia was removed by vacuum concentration. The oligonucleotide pool was purified using 2 x illustra NAP-5 Columns (GE Life Sciences, PN: 17085302). The resulting solution was evaporated to dryness, resuspended in water and quantified by Nanodrop absorbance at 260 nm. We used ~0.1 fmol of each individual oligonucleotide per enrichment reaction.

We ordered panels of double-stranded capture oligonucleotides from Twist Bioscience (Custom probes for NGS target enrichment). As recommended by Twist, we used 13.67 fmol of each individual oligonucleotide per enrichment reaction.

#### Experimental procedure

For samples containing 100,000 cells or fewer, we followed a low-input 3C library preparation protocol^49^. Cells were sorted into 1 ml medium and fixed with 2% formaldehyde for 10 minutes. After the reaction was quenched with glycine, cells were pelleted and washed with cold PBS. Following centrifugation, ~5% supernatant was left behind to avoid disturbing the pellet. Cells were resuspended in cold lysis buffer (10 mM Tris pH 8, 10 mM NaCl, 0.2% Igepal CA-630, 1x cOmplete Protease Inhibitor Cocktail, water) and snap frozen. Prior to digestion, cells were pelleted and all lysis buffer was removed by careful pipetting. Chromatin was subsequently digested with the DpnII restriction enzyme in a 200 μl reaction, to which 3 doses of 150 U DpnII enzyme were added several hours apart and which was incubated 16–24 hours at 37 °C. After heat-inactivation of DpnII, the ligation reaction was performed in the same tube, using 120 U T4 ligase in an overnight incubation at 16 °C. Ligated DNA was reverse crosslinked and treated with proteinase K at 65 °C in an overnight incubation. After RNase treatment, DNA was extracted using phenol-chloroform and transferred to a light phase lock tube for separation. To maximize yield, DNA was precipitated overnight in 70% ethanol at −20 ºC, after which DNA was pelleted and resuspended in PCR-grade water. The resulting 3C libraries were sonicated to 200 bp fragments using a Covaris S220 Focused Ultrasonicator (six cycles of 60s; duty cycle: 10%; intensity: 5; cycles per burst: 200). Illumina TruSeq adapters were subsequently added using the NEBNext Ultra II DNA Library Prep kit, according to the manufacturer’s protocol. DNA clean up steps were performed with Ampure XP beads at a 1:1.8 ratio to minimize loss of material. The libraries were indexed and amplified in 8-12 rounds of PCR amplification using the Herculase II PCR kit. 700 ng–1 μg of indexed material was used during subsequent capture-based enrichment.

When cell numbers were not limiting, we used aliquots of ~10^7^ cells to prepare 3C libraries, following the efficient Capture-C protocol ^22^. Cells were fixed with 2% formaldehyde for 10 minutes, after which the reaction was quenched with glycine. The fixed cells were washed in cold PBS and resuspended in cold lysis buffer. After incubation on ice for 20 minutes, the cells were snap frozen. Prior to digestion, the cells were resuspended in restriction buffer, Dounce homogenized on ice, and treated with SDS and Triton X-100. The chromatin was digested with DpnII, using 3 aliquots of 1500 U DpnII restriction enzyme, which were added several hours apart over a total incubation time of 16–24 hours at 37 °C. The digestion reaction was heat-inactivated and digested chromatin was ligated overnight with 720 U of T4 DNA ligase at 16 °C. The ligated DNA was reverse crosslinked and treated with proteinase K overnight at 65 °C. After RNase treatment, DNA was purified using phenol-chloroform extraction and precipitation with ethanol and sodium acetate at −80 °C. The resulting 3C libraries were resuspended in PCR-grade water. Aliquots of 5–6 μg of 3C library were sonicated to ~200 bp fragments using a Covaris S220 Focused Ultrasonicator (six cycles of 60s; duty cycle: 10%; intensity: 5; cycles per burst: 200). Illumina TruSeq adaptors were added using NEBNext DNA Library Prep reagents according to the manufacturer’s protocol. The libraries were indexed and amplified using Agilent Herculase II reagents in a 6-cycle PCR reaction. DNA clean-up steps were performed using AMPure XP beads in a 1.8:1 bead:sample ratio. Where possible, we processed 2 aliquots of each sample in parallel and ligated the DNA with the same index to generate maximum library complexity. 1–1.5 μg of indexed material was used during subsequent capture-based enrichment.

For enrichment using single-stranded oligonucleotides, we used the Nimblegen SeqCap EZ reagents and followed the SeqCap EZ Library SR User’s Guide (Chapters 5–7). We multiplexed up to 6 samples per enrichment reaction in a single tube, and multiplied the volumes described in the protocol by the number of multiplexed libraries. Briefly, 700 ng–1.5 μg of indexed library (in exact 1:1 ratio if multiplexed reaction is performed) was mixed with 5 μg mouse Cot-1 DNA, 1 nmol of Nimblegen HE universal blocking oligonucleotides and 1 nmol of Nimblegen HE index-specific blocking oligonucleotides (corresponding to the Illumina TS index used) in a 1.5 μl microcentrifuge tube. This mixture was dried completely in a vacuum centrifuge at 50°C. The dried DNA was resuspended in 7.5 μl Nimblegen hybridization buffer and 3 μl Nimblegen hybridization component A and denatured at 95°C for 10 minutes. Concurrently, 4.5 μl of the biotinylated capture oligonucleotides was placed in a 200 μl PCR tube and heated to 47 °C in a PCR thermocycler. After denaturation, the 3C library mixture was added to the biotinylated oligonucleotides without removing them from the heating block in the thermocycler. The hybridization reaction was incubated in the thermocycler at 47 °C for 64–72 h with a heated lid at 57 °C. After incubation was complete, we enriched for the captured DNA fragments using M270 streptavidin beads and the Nimblegen SeqCap EZ wash buffers, following the procedure described in the manufacturer’s protocol. After the washing steps, the beads with captured material were resuspended in 40 μl PCR-grade water. Captured DNA was amplified directly of the beads using the KAPA master mix provided in the SeqCap EZ accessory kit v2, in 2 separate reactions of ~10 cycles, as described in the protocol. Ampure-XP beads were used in a 1.8:1 bead:sample ratio to clean up the amplification reaction and DNA was eluted in 30 μl PCR-grade water. To increase enrichment, a second round of oligonucleotide capture was performed following the same procedure, using up to 2 ug of enriched material in a single hybridization reaction (even if multiplexed in first round) of 20–24 hours. For enrichment using double-stranded oligonucleotides, we used Twist Biosciences reagents and followed the Twist Custom Panel Protocol (Steps 4–7). To multiplex samples, we used 375–500 ng indexed library per sample, mixed up to 1.5 μg (in exact 1:1 ratio) in a single tube, and used single reaction volumes as described in the protocol. We processed multiple tubes simultaneously if required. Streptavidin C1 beads were used to enrich the hybridized DNA and the washed material was amplified using 10–12 cycles of PCR. Ampure-XP beads were used in a 1.8:1 bead:sample ratio to clean up the amplification reaction and DNA was eluted in 30 μl PCR-grade water. To increase enrichment, a second round of oligonucleotide capture was performed following the same procedure, using up to 1.5 μg of enriched material in a single hybridization reaction (even if multiplexed in first round) of 20–24 hours. The enriched Tiled-C libraries were assessed using the Agilent Bioanalyzer or D1000 Tapestation and quantified using KAPA Library Quantification reagents, before sequencing using the Illumina NextSeq platform. In high-quality libraries, sequencing 3–5 million reads per enriched Mb per sample is sufficient for data at 5 kb resolution.

#### Analysis

The most straightforward way to analyze Tiled-C data is to use the HiCPro pipeline^50^ with the options for Capture-Hi-C analysis. We have also adjusted our pipelines for Capture-C analysis to be compatible with Tiled-C data. This pipeline is designed to analyze deep, targeted 3C data and provides very stringent filtering, especially regarding PCR-related artefacts. All data presented in the paper have been analyzed using a combination of this CCseqBasic pipeline (https://github.com/Hughes-Genome-Group/CCseqBasicF/releases), custom scripts and ICE normalization^51^. Detailed instructions and custom scripts are available on https://github.com/oudelaar/TiledC. Interaction profiles from virtual viewpoints (Supplementary Figure 1) can be generated using the CCseqBasic pipeline with the options for Capture-C analysis.

When Tiled-C data are compared between different cell types (Figures 1b, 1c, 4, 5; Supplementary Figures 2, 7-12), we have down-sampled the data in the different samples to an equal number of valid interactions within the tiled region.

Quantification of enhancer-promoter interactions of interest was performed based on interaction counts in the corresponding bins after normalizing for the total number of counts in the matrix. We used the following coordinates for quantification of regulatory elements of interest (highlighted in Supplementary Figures 10-12): α-globin: chr11:32,145,000-32,148,000; chr11:32,182,000-32,185,000; chr11:32,195,000-32,198,000; Slc25a37: chr14:69,902,000-69,905,000; chr14:69,939,000-69,942,000; Tal1: chr4:114,728,000-114,731,000; chr4:114,768,000-114,771,000; Cd47: chr16:49,770,000-49,773,000 and chr16:49,852,000-49,855,000; Cpeb4: chr11:31,711,000-31,714,000; chr11:31,770,000-31,773,000; Btg: chr1:135,995,000-135,998,000; chr1:135,975,000-135,978,000. We identified TADs based on insulation indices using TADtool^52^.

To examine the reproducibility of Tiled-C in low-input samples, we used HiCRep^53^ to calculate stratum-adjusted correlation coefficients, considering a maximum distance of 100,000 bp.

### Hi-C

We compared Tiled-C data in mouse ES cells to the deepest currently available Hi-C data in mouse ES cells^9^. We explored the data using HiGlass^54^ and downloaded and re-analyzed the Hi-C data using the HiC-Pro pipeline^50^ with default options and ICE normalization^51^.

### ATAC-seq

#### Experimental procedure

For FACS-sorted erythroid progenitors from fetal liver, either 2 or 3 replicates were processed of ~50,000 cells each for each sorted population. ATAC-seq was performed as previously described^35^.

For FACS-sorted hematopoietic stem and progenitor cells from adult bone marrow, 1 replicate of between 5,000 and 20,000 cells was processed for each population. Cells were spun at 500 g for 10 minutes at 4 °C. The supernatant was discarded and cells were resuspended directly in Dig-transposition buffer (25 μl 2x TD Buffer [Illumina], 2.5 μl Tn5 transposase, 0.5 μl 1% digitonin and 22 μl H_2_O) before incubating at 37 °C for 30 minutes with agitation at 600 rpm. After the transposition step, samples were processed as previously described^35^.

#### Analysis

Reads were mapped to the mouse mm9 genome and PCR duplicates removed using NGseqBasic^55^. Technical replicates were merged and peaks called using MACS2^56^. Peaks were merged and the number of reads in each sample overlapping each peak was calculated using BEDTools merge and multicov^57^. For visualization, bedgraph files were generated using BEDTools genomecov with a scaling factor of 1e6 / (total number of reads in peaks). All analysis scripts are available at https://github.com/rbeagrie/alpha-tiledc.

### Single-cell RNA-seq

#### Experimental procedure

Fetal livers were harvested and pooled from 7 e13.5 C57BL/6 mouse embryos and processed as above. Cells were first stained with 2.5 μg/ml biotin-conjugated anti-Ter119 (BD 553672) and 2.5 μg/ml purified anti-Ter119 (BioLegend 116241) conjugated to a DNA oligonucleotide (ADT2 GAGGCGATTGAT). After magnetic depletion, an equal number of Ter119+ cells were added back to the Ter119-fraction for further analysis, ensuring a balanced distribution of erythroid cells from all stages of differentiation. Cells were then stained with a panel of 9 further DNA-oligonucleotide conjugated antibodies (0.33 μg/ml anti-CD71: BioLegend 113802 conjugated with ADT9 [CGAAGAAGGAGT], 1 μg/ml anti-CD41: BioLegend 133919 conjugated with ADT3 [TGTCCGGCAATA], 1 μg/ml anti-CD45R: BioLegend 103249 conjugated with ADT5 [GATCGTAATACC], 1 μg/ml anti-CD3e: BioLegend 100345 conjugated with ADT7 [CATCGGTGTACA], 1 μg/ml anti CD11b: BioLegend 101249 conjugated with ADT1 [CATGATTGGCTC], 1 μg/ml anti-Ly6G/6C: BioLegend 108449 conjugated with ADT4 [TGGTGAACCTGG], 1 μg/ml anti-CD44: BioLegend 103051 conjugated with ADT8 [GTCTAGACTTCG], 1 μg/ml anti-cKit: BioLegend 105829 conjugated with ADT6 [AAGCGCTTGGCA], and 1 μg/ml non-specific IgG conjugated with ADT10 [CGGAGTAGTAAT]). Antibodies were conjugated to streptavidin as previously described^34^ and mixed with biotinylated custom oligonucleotides. Cells were processed for single-cell RNA-seq using the 10x Genomics Single Cell 3’ v2 kit.

#### Analysis

cDNA reads were mapped to the mouse mm10 assembly and barcodes assigned to cells using Cell Ranger v2.1.1 (10x Genomics). ADT reads were mapped to cell/antibody barcodes using CITE-seq-count (https://github.com/Hoohm/CITE-seq-Count). Potential doublet cells were removed using Scrublet^58^. Further analysis was performed using Seurat v2. Low quality cells with less than 300 or more than 5,000 identified genes, or with more than 9% mitochondrial reads were also removed. Clusters were identified using the “FindClusters” function and UMAP projection was generated using the “RunUMAP” function, both with the first 16 principle components. Seurat clusters were annotated using marker genes and by reference to previously published data^33^ – Seurat identified two clusters corresponding to committed erythroid progenitors (CEP) and four clusters corresponding to cells undergoing erythroid terminal differentiation (ETD). Average gene expression for populations matching those obtained by FACS sorting was generated by using the “SubsetData” function to select cells with low levels of cell-surface barcodes corresponding to lineage markers, and appropriate levels of barcodes corresponding to CD71 and Ter119. All analysis scripts are available at https://github.com/rbeagrie/alpha-tiledc.

### RNA-FISH

#### Experimental procedure

Standard RNA-FISH was carried out as previously described^59^. Sorted cells from mouse fetal liver were placed back into culture for 6 hours to allow nascent transcription to be re-established. Samples were hybridized with digoxygenin-labelled oligonucleotide probes directed to α-globin introns (30 ng per slide) and visualized using FITC-conjugated antibodies (primary: sheep anti-DIG FITC [Roche] 1:50, secondary: rabbit anti-sheep FITC [Vector] 1:100). Two negative controls were also included: brain tissue from a male, adult CD1 mouse and Ter119+ (i.e. mature) fetal liver erythroid cells that were probed with secondary antibody but no primary antibody. Magnetically purified but not FACS-purified Ter119+ fetal liver erythroid cells were used as a positive control.

#### Imaging equipment and settings

Widefield fluorescence imaging was performed at 20 °C on a DeltaVision Elite system (Applied Precision) equipped with a 100 ×/1.40 NA UPLSAPO oil immersion objective (Olympus), a CoolSnap HQ2 CCD camera (Photometrics), DAPI (excitation 390/18; emission 435/40) and FITC (excitation 475/28; emission 525/45) filters. 12-bit image stacks were acquired with a z-step of 150 nm giving a voxel size of 64.5 nm × 64.5 nm × 150 nm.

#### Image analysis

Image analysis was blinded by renaming image files from all experiments with random character strings and processing them together. Images were manually examined, and each cell was scored for the presence of active nascent-transcription foci. Analysis scripts are available at https://github.com/rbeagrie/alpha-tiledc.

## Data availability

Tiled-C, ATAC-seq and single-cell RNA-seq data generated in this study have been deposited in the Gene Expression Omnibus (GEO) under accession code GSE137477. All RNA-FISH image files are archived in Dryad. A UCSC hub for visualizing ATAC-seq and single-cell RNA-seq mean expression data is available at http://sara.molbiol.ox.ac.uk/public/hugheslab/alpha-tiledc/hub.txt.

## Acknowledgements

We thank J. Sloane-Stanley and J. Sharpe of the WIMM Transgenics Facility for mouse breeding and fetal liver provision, K. Clark, S-A. Clark and C. Waugh of the WIMM Flow Cytometry Facility for cell sorting services, N. Ashley of the WIMM Single Cell facility for antibody conjugation and 10X library preparation services, J. Brown for help with RNA-FISH, D. Downes for help with ATAC-seq and C. Garnett for advice on low-input ATAC-seq. We wish to acknowledge the Centre for Computational Biology, MRC Weatherall Institute of Molecular Medicine, Radcliffe Department of Medicine and University of Oxford for use of their services in this project. This work was supported by the Medical Research Council (MC_UU_00016 to D.R.H., J.R.H., V.J.B. and MR/N00969X/1 to J.R.H., T.B., V.J.B.), Wellcome Trust (209181/Z/17/Z to R.A.B.), National Institutes of Health (NIH NIDDK R01DK100915 to M.S.), Biotechnology and Biological Sciences Research Council (BB/J001694/2 and BB/ R008655/1 to A.H.E.-S. and BB/M025624/1 to A.S.) and a Stevenson Junior Research Fellowship at University College, Oxford (to A.M.O.).

## Author Contributions

A.M.O. and R.A.B. performed experiments and bioinformatic analyses and wrote the manuscript. M.G., E.G., D.H. and J.C. performed experiments. S.d.O., A.S., A.H.E.-S. and T.B. synthesized oligonucleotides. J.K. wrote the oligonucleotide design tool. J.M.T. performed bioinformatic analyses. V.J.B. and M.S. provided conceptual advice. D.R.H. and J.R.H. supervised the project and wrote the manuscript.

