## Supplementary Figures and Tables for "Dynamics of the 4D genome during lineage specification, differentiation and maturation *in vivo*"

**Supplementary Figure 1: Tiled-C contact matrix and virtual viewpoints of the α-globin locus.**

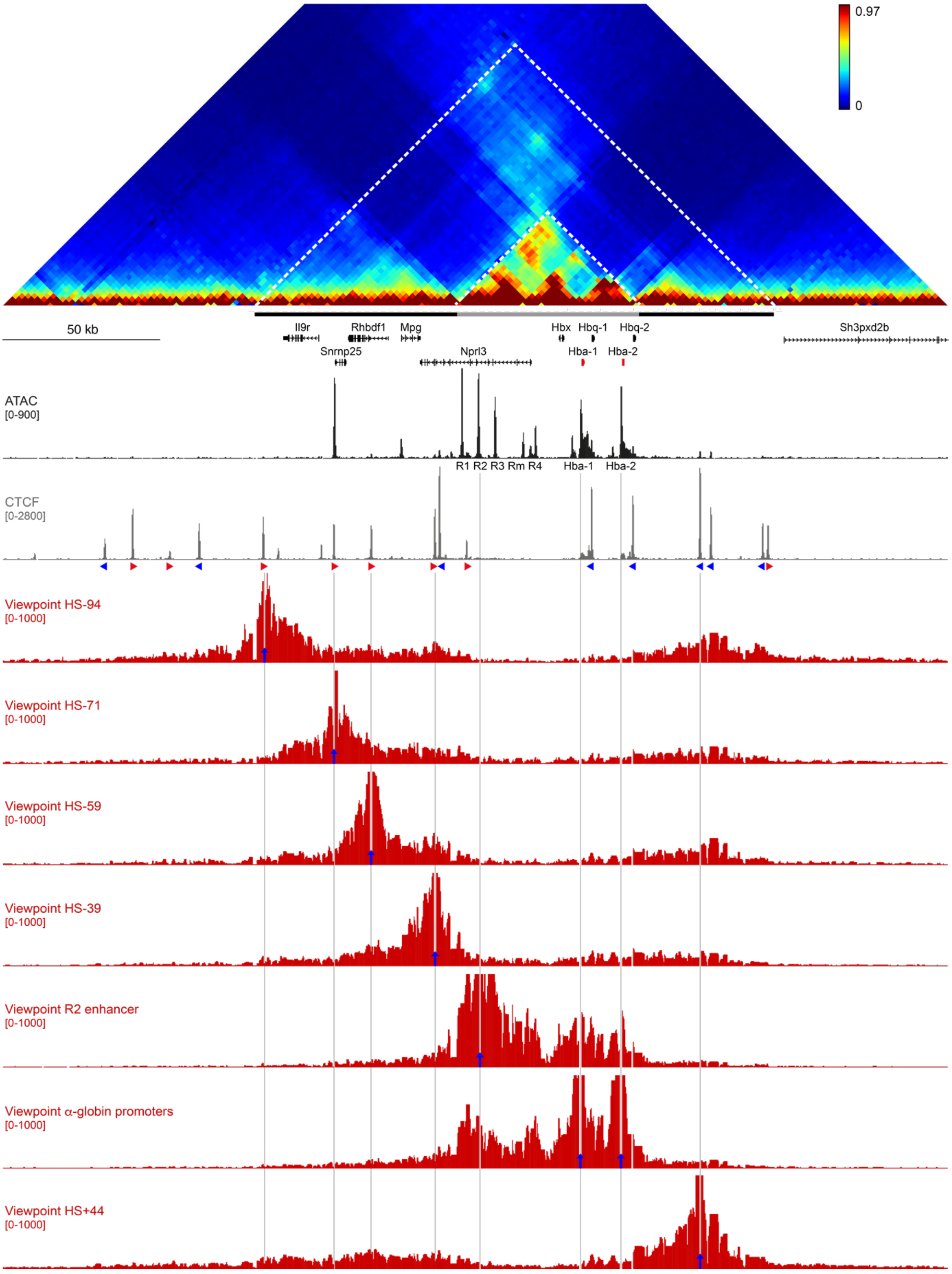

**Supplementary Figure 1: Tiled-C contact matrix and virtual viewpoints of the $\alpha$-globin locus. [continued]**

Tiled-C contact matrix of 300 kb spanning the $\alpha$-globin locus in primary mouse erythroid cells at 2 kb resolution. The pre-existing TAD and sub-compartmentalized erythroid-specific domain are indicated below the matrix with a black and grey bar and highlighted in the matrix with white dashed lines. Gene annotation is shown underneath, with the $\alpha$-globin genes highlighted in red. Open chromatin (ATAC) is shown below, with the $\alpha$-globin enhancers R1-R4 and Rm indicated. CTCF occupancy is shown underneath, with the orientation of the CTCF-binding motifs indicated by arrowheads (forward orientation in red; reverse orientation in blue). Interaction profiles from virtual viewpoints (highlighted by blue arrows) across the $\alpha$-globin locus are shown at the bottom. Contact frequencies represent normalized, unique interactions in 3 replicates. Coordinates (mm9): chr11:32,000,000-32,300,000.

**Supplementary Figure 2: Tiled-C contact matrices of the α-globin locus.**

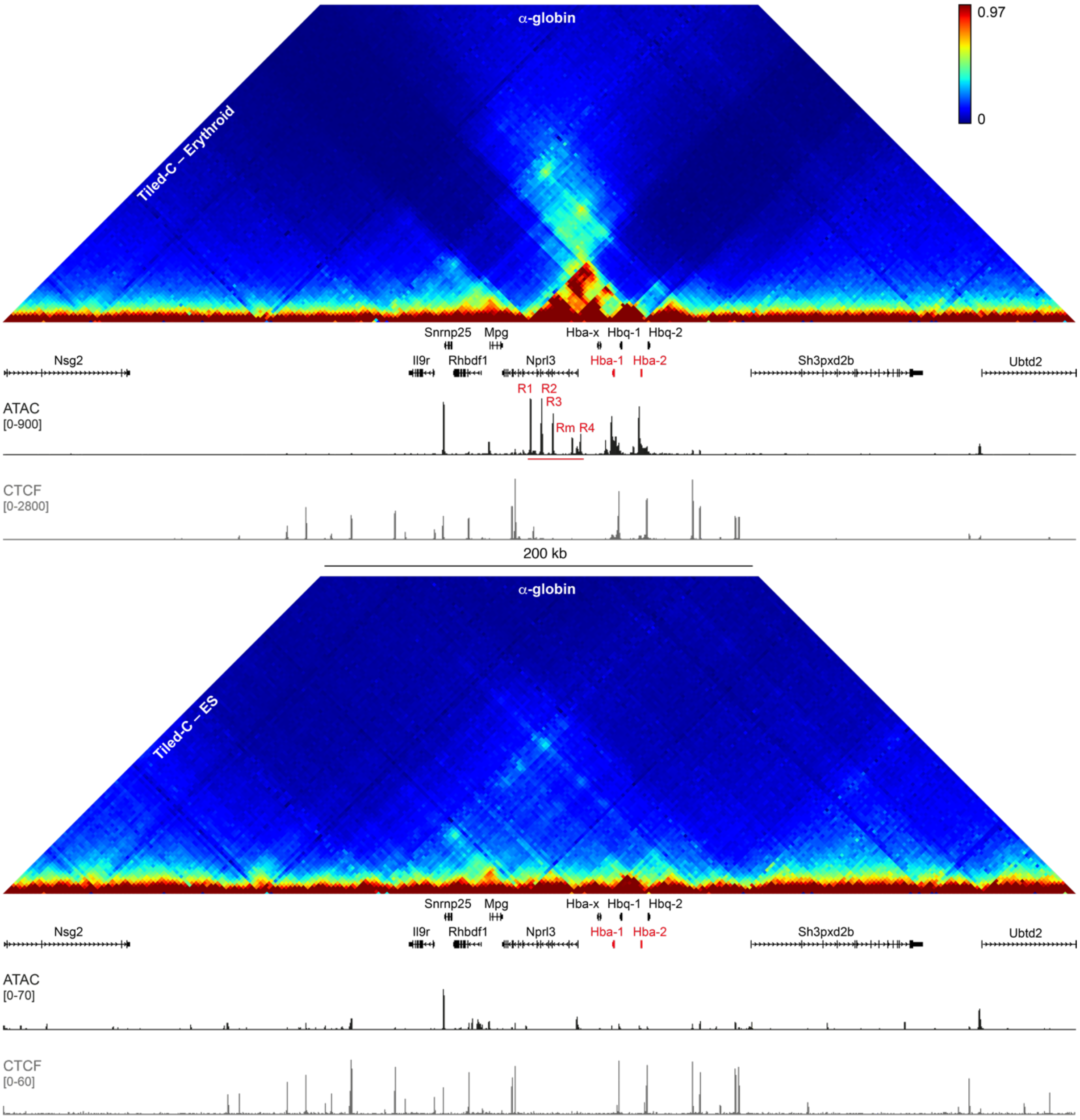


Tiled-C contact matrices of 500 kb spanning the α-globin locus in primary mouse erythroid cells (top) and ES cells (bottom) at 2 kb resolution. Contact frequencies represent normalized, unique interactions in 3 replicates. Gene annotation (α-globin genes highlighted in red), open chromatin (ATAC; α-globin enhancers highlighted in red) and CTCF occupancy are shown below the matrices. Coordinates (mm9): chr11:31,900,000-32,400,000.

**Supplementary Figure 3: Reproducibility of Tiled-C data derived from low-input samples.**

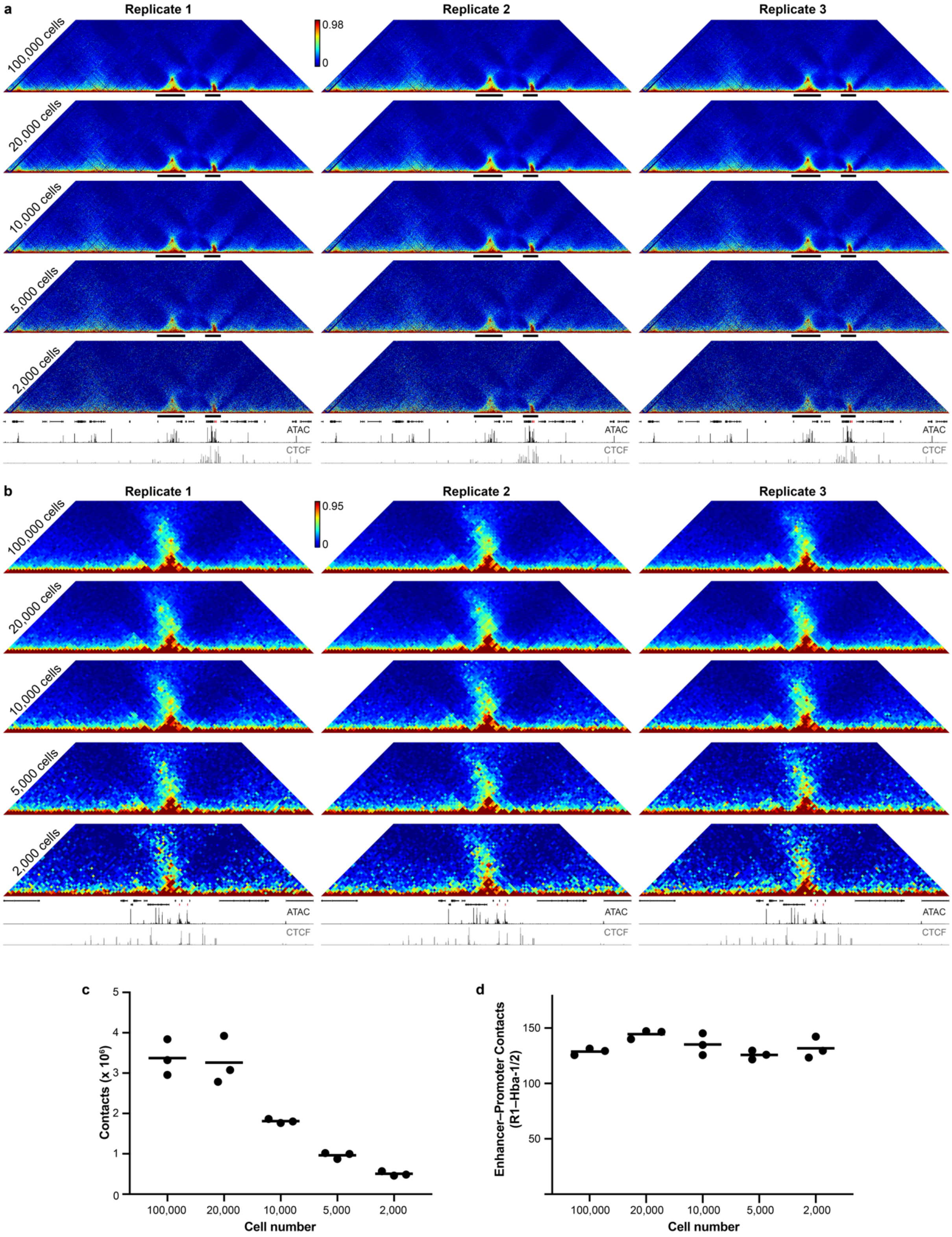

**Supplementary Figure 3: Reproducibility of Tiled-C data derived from low-input samples. [continued]**

**(a)** Tiled-C contact matrices of ~3.3 Mb spanning the α-globin locus in individual biological replicates of small aliquots of primary mouse erythroid cells at 5 kb resolution. Contact frequencies represent normalized, unique interactions. TADs are indicated below the matrix with a black bar. Gene annotation (α-globin genes highlighted in red), open chromatin (ATAC) and CTCF occupancy are shown below the matrices. Coordinates (mm9): chr11:29,900,000-33,230,000. Stratum-adjusted correlation coefficients between individual replicates are 0.979-0.983 for 100,000 cells, 0.987-0.990 for 20,000 cells, 0.972-0.980 for 10,000 cells, 0.967-0.971 for 5,000 cells, and 0.953-0.962 for 2,000 cells. **(b)** Tiled-C contact matrices of 500 kb spanning the α-globin locus in individual biological replicates of small aliquots of primary mouse erythroid cells at 5 kb resolution. Contact frequencies represent normalized, unique interactions. Gene annotation (α-globin genes highlighted in red), open chromatin (ATAC) and CTCF occupancy are shown below the matrices. Coordinates (mm9): chr11:31,900,000-32,400,000. **(c)** Number of total, unique contacts in the α-globin locus (chr11:29,900,000-33,230,000) identified in individual replicates of small aliquots of primary erythroid cells. At this sequencing depth (Supplementary Table 2), library complexity is limiting when fewer than 20,000 cells are used. **(d)** Normalized contact frequencies between the α-globin promoters and the R1 enhancer in individual replicates of small aliquots of primary erythroid cells, indicating that enhancer-promoter interactions are reliably detected across individual replicates.

**Supplementary Figure 4: Tiled-C data generated with single- and double-stranded capture oligonucleotides.**

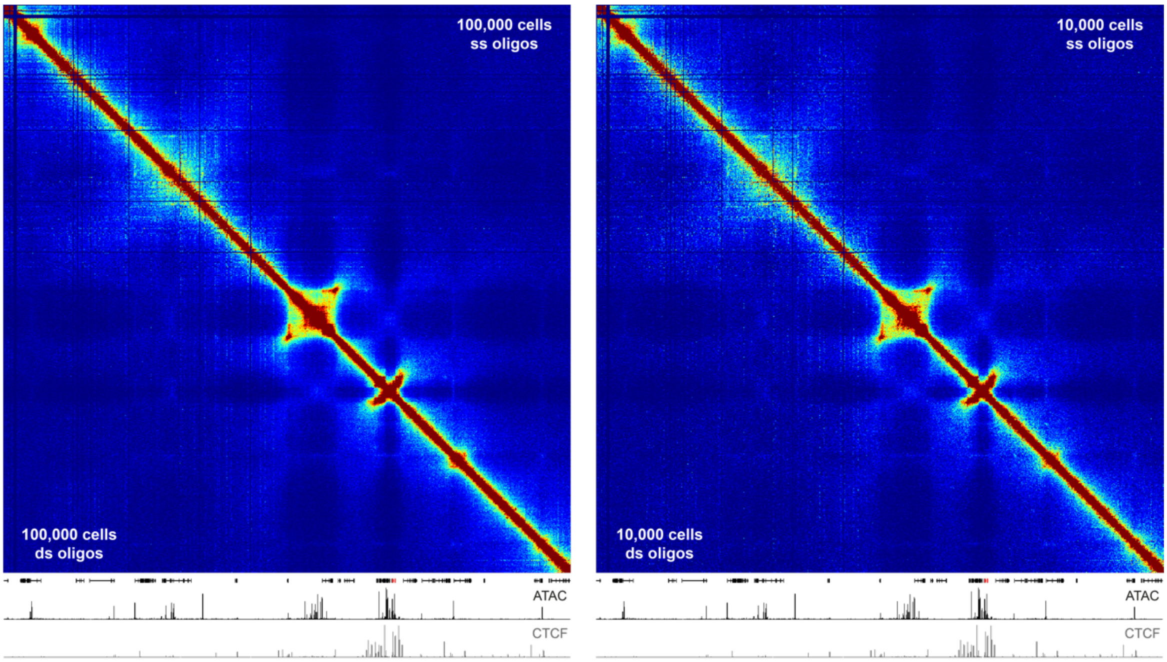


Tiled-C contact matrices of 3.3 Mb spanning the α-globin locus at 5 kb resolution generated using single-stranded (ss; top) and double-stranded (ds; bottom) capture oligonucleotides in aliquots of 100,000 (left) and 10,000 (right) primary erythroid cells. Contact frequencies represent normalized, unique interactions in 3 replicates. Gene annotation (α-globin genes highlighted in red), open chromatin (ATAC) and CTCF occupancy are shown below the matrices. Coordinates (mm9): chr11:29,900,000-33,230,000.
The contact matrices are nearly identical, indicating that the use of single or double-stranded capture oligonucleotides does not introduce any technical variability.

### Supplementary Figure 5: Single-cell RNA-seq.

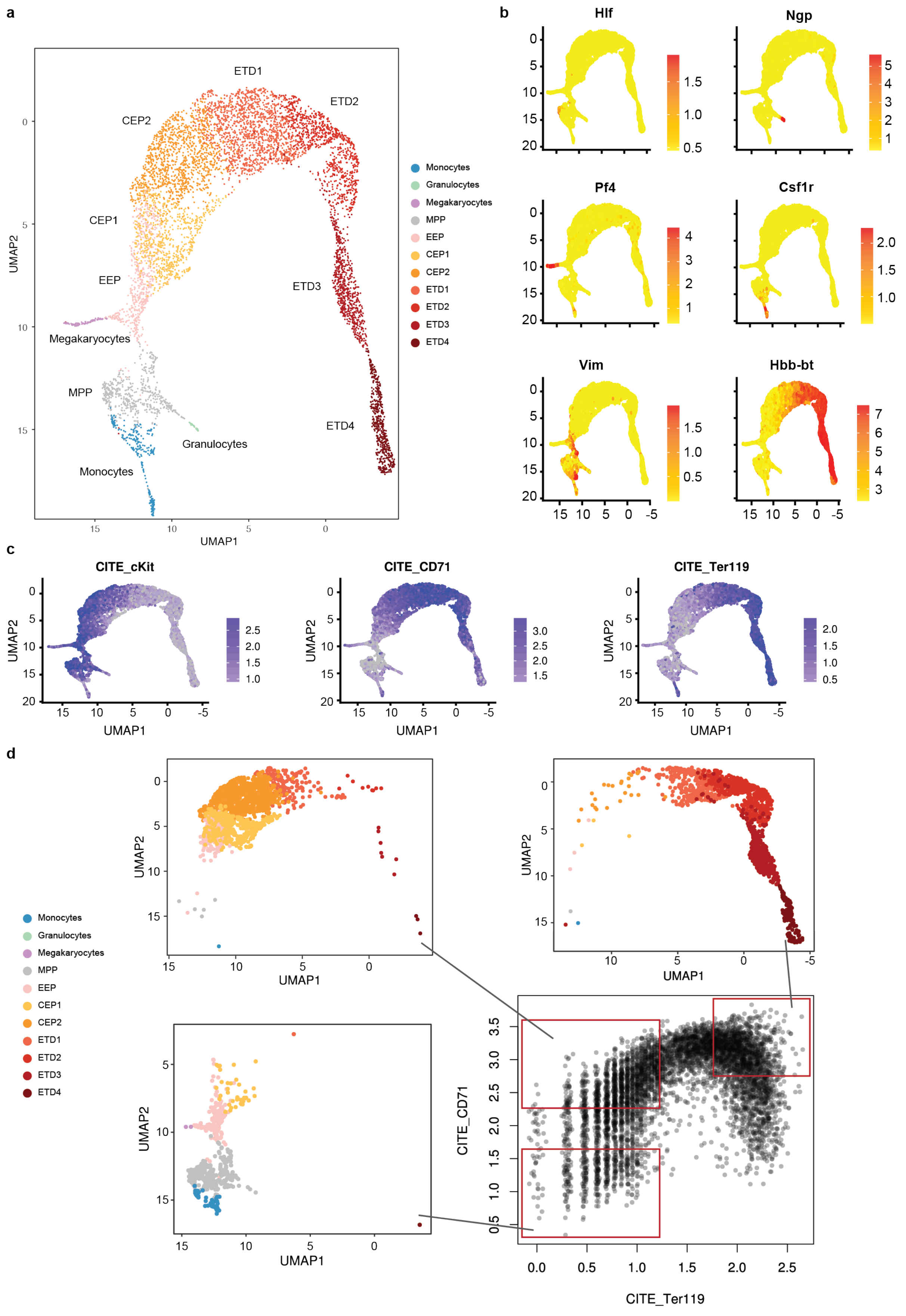

**Supplementary Figure 5: Single-cell RNA-seq. [continued]**

**(a)** UMAP projection of Cellular Indexing of Transcriptomes and Epitopes by Sequencing (CITE-seq) data showing cell clusters detected by Seurat. Based on marker gene expression, three smaller trajectories were assigned as Monocytes, Granulocytes and Megakaryocytes, whilst cells in the center of these trajectories were identified as multi-potent progenitors (MPP). Based on previously published scRNA-seq data (Tusi et al. Nature 2018), the erythroid clusters were identified as early erythroid progenitors (EEP), committed erythroid progenitors (CEP1-2) or cells undergoing erythroid terminal differentiation (ETD1-4). **(b)** Example marker genes used to annotate scRNA-seq clusters, displayed with Seurat normalized expression values. **(c)** Cells were also labelled with CITE-seq barcoded antibodies against the same surface markers used for FACS purification. Plots show three example surface markers highlighting early populations (cKit), mid-differentiation (CD71) and late-differentiation (Ter119) erythroid cells. **(d)** *In silico* gating of the scRNA-seq data was used to match cell populations from scRNA-seq to bulk datasets obtained through FACS sorting. *Bottom right:* Gating of Lin- cells based on barcode counts for CD71 and Ter119. *Bottom left, top left, top right:* For three example gates (S0-low, S1 and S3, respectively) the cluster composition of the included cells is shown. S0-low contains predominantly MMP and EEP, S1 contains mostly CEP and S3 contains ETD cells.

**Supplementary Figure 6: Chromatin accessibility in the α-globin locus through erythroid differentiation.**

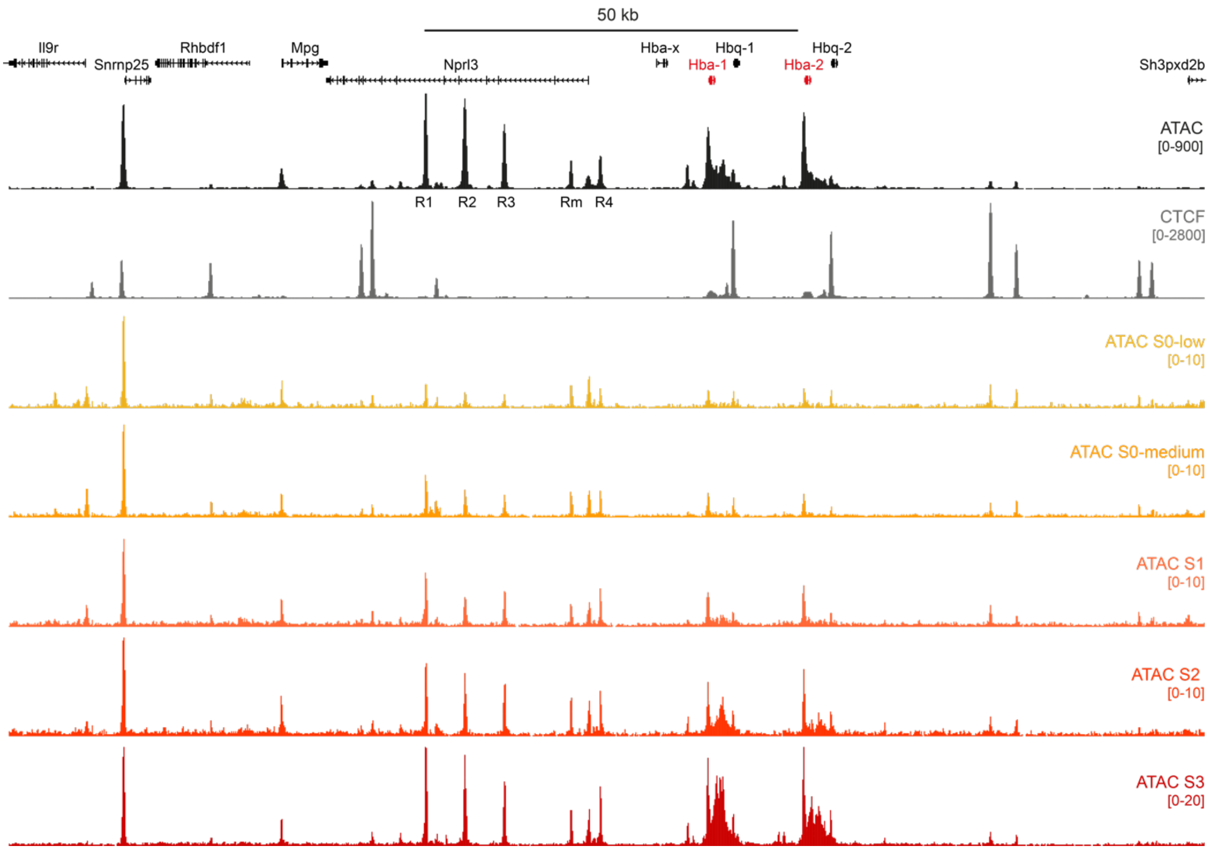


Chromatin accessibility (ATAC) in a region of 160 kb spanning the mouse α-globin locus in sequential stages of *in vivo* erythroid differentiation. ATAC profiles represent normalized data from 3 S0-low, S0-medium and S1 replicates and 2 S2 and S3 replicates. The profiles are shown at different scales to highlight changes in accessibility in early stages of differentiation. Gene annotation (α-globin genes highlighted in red), open chromatin (ATAC; α-globin enhancers R1-R4 and Rm indicated) and CTCF occupancy in mature mouse erythroblast cells are shown at the top. Coordinates (mm9): chr11:32,090,000-32,250,000.

**Supplementary Figure 7: Chromatin interactions and accessibility in the extended α-globin locus through erythroid differentiation.**

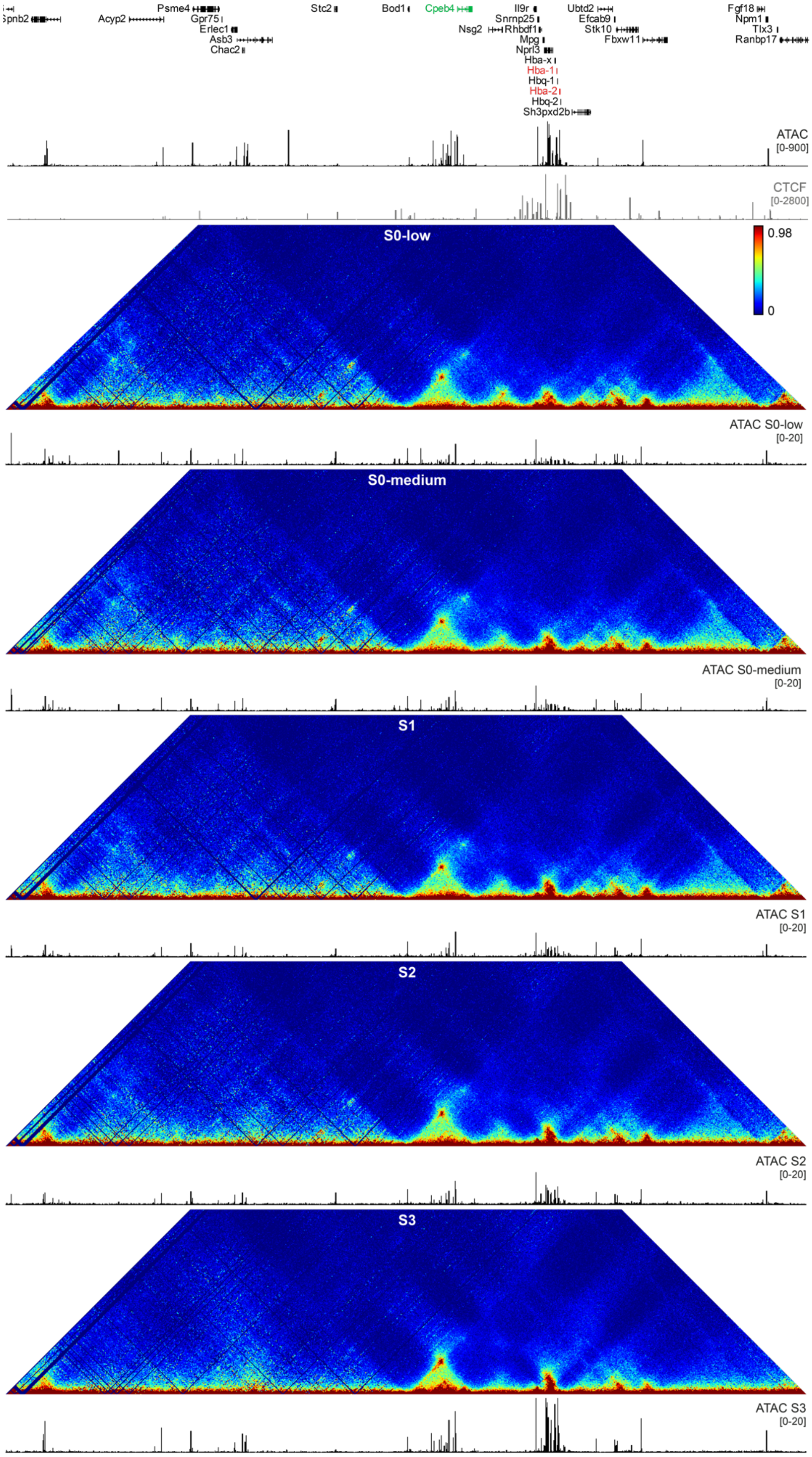

**Supplementary Figure 7: Chromatin interactions and accessibility in the extended α-globin locus through erythroid differentiation. [continued]**

Tiled-C contact matrices of ~3.3 Mb spanning the mouse α-globin locus in sequential stages of *in vivo* erythroid differentiation at 5 kb resolution. Contact frequencies represent normalized, unique interactions in 2 replicates. Matched open chromatin (ATAC) profiles are shown underneath each matrix and represent normalized data from 3 S0-low, S0-medium and S1 replicates and 2 S2 and S3 replicates. Gene annotation (α-globin and *Cpeb4* genes highlighted in red and green, respectively), open chromatin (ATAC) and CTCF occupancy in mature mouse erythroblast cells are shown at the top. Coordinates (mm9): chr11:29,900,000-32,230,000.

**Supplementary Figure 8: Reproducibility of chromatin interactions in the α-globin locus through erythroid differentiation.**

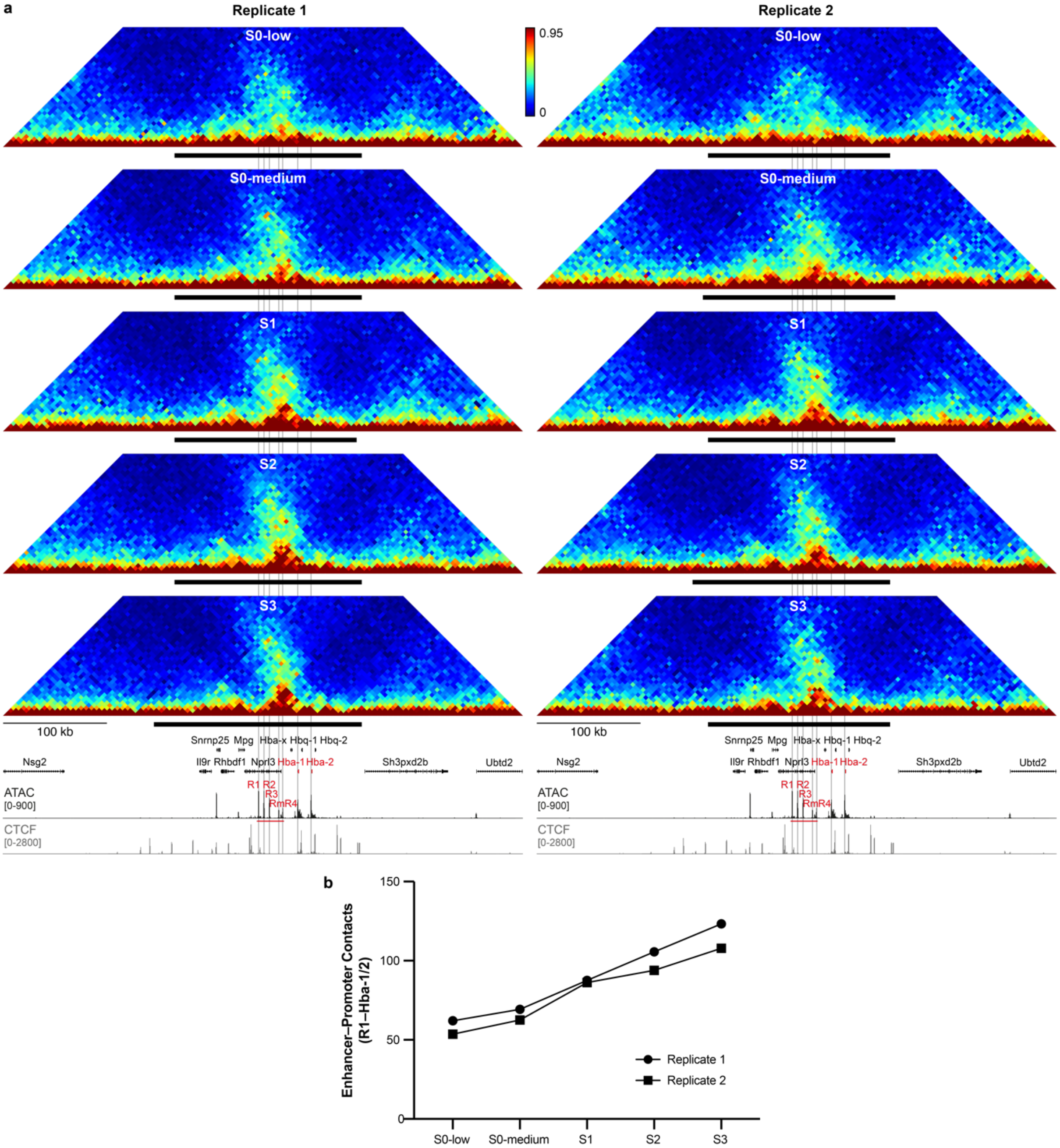


**(a)** Tiled-C contact matrices of 500 kb spanning the mouse α-globin locus in individual biological replicates of sequential stages of *in vivo* erythroid differentiation at 5 kb resolution. Contact frequencies represent normalized, unique interactions. TADs are indicated below each matrix with a black bar. Gene annotation (α-globin genes highlighted in red), open chromatin (ATAC; α-globin enhancers indicated in red) and CTCF occupancy are shown underneath the matrices. Coordinates (mm9): chr11:31,900,000-32,400,000. **(b)** Normalized contact frequencies between the α-globin promoters and the R1 enhancer in individual biological replicates of sequential stages of *in vivo* erythroid differentiation.

**Supplementary Figure 9: Chromatin interactions and accessibility in the α-globin locus in early hematopoietic progenitor populations.**

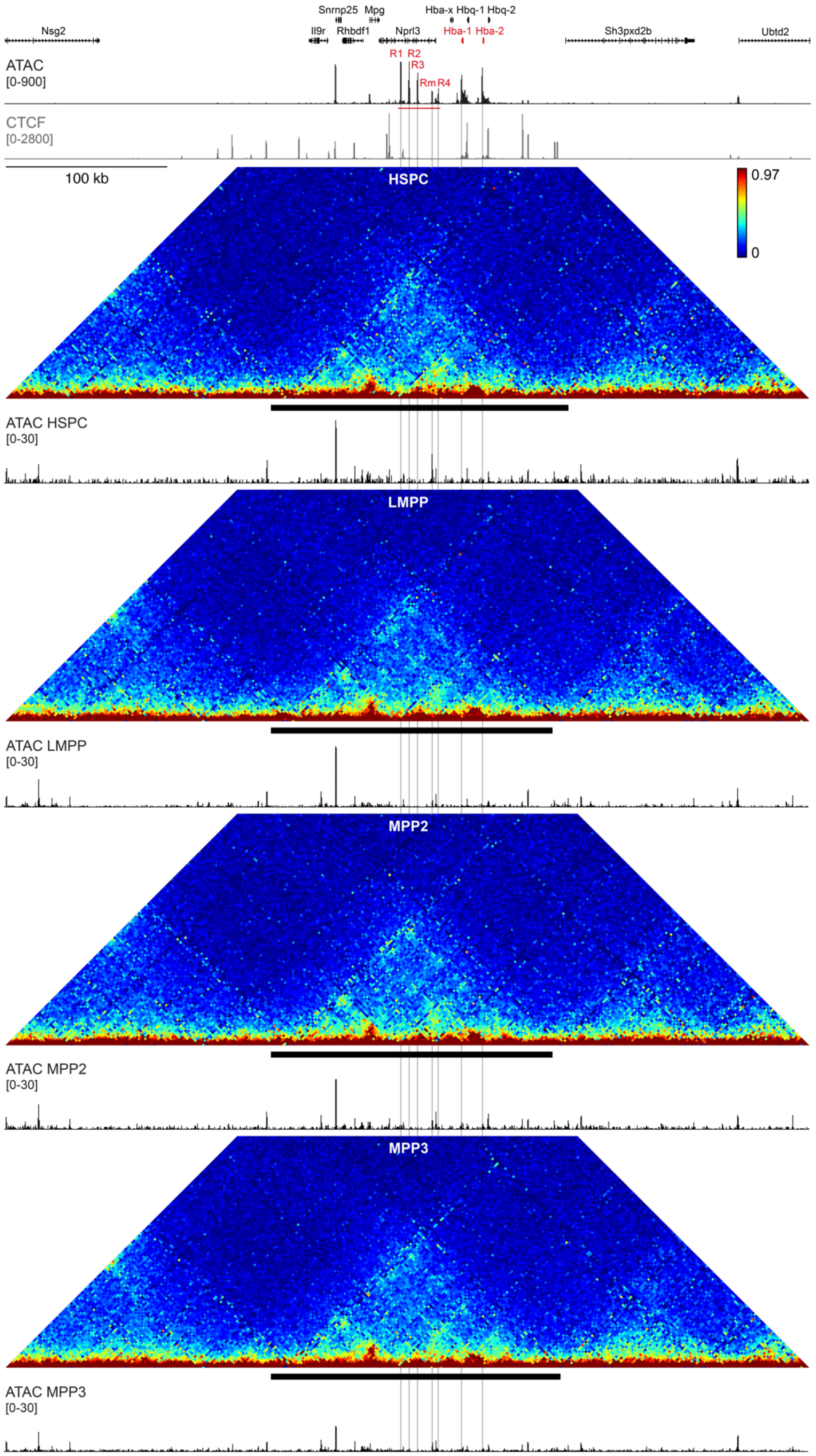

**Supplementary Figure 9: Chromatin interactions and accessibility in the α-globin locus in early hematopoietic progenitor populations. [continued]**

Tiled-C contact matrices of 500 kb spanning the mouse α-globin locus in early hematopoietic progenitor populations at 2 kb resolution. Contact frequencies represent normalized, unique interactions from 2 replicates. TADs are indicated below the matrices with a black bar and matched open chromatin (ATAC) profiles are shown underneath. Gene annotation (α-globin genes highlighted in red), open chromatin (ATAC; α-globin enhancers indicated in red) and CTCF occupancy in mature mouse erythroblast cells are shown at the top. Coordinates (mm9): chr11:31,900,000-32,400,000.
Hematopoietic stem and progenitor cells (HSPCs) differentiate into multipotent progenitors (MPPs), which can give rise to both the myeloid lineage – from which erythroid cells are formed – and the lymphoid lineage through the formation of lymphoid-primed multipotential progenitors (LMPPs). With exception of low levels of accessibility for the Rm enhancer, the α-globin enhancers are not accessible in the early hematopoietic progenitor populations.

**Supplementary Figure 10: Chromatin interactions and accessibility in the *Slc25a37* and *Tal1* loci through erythroid differentiation.**

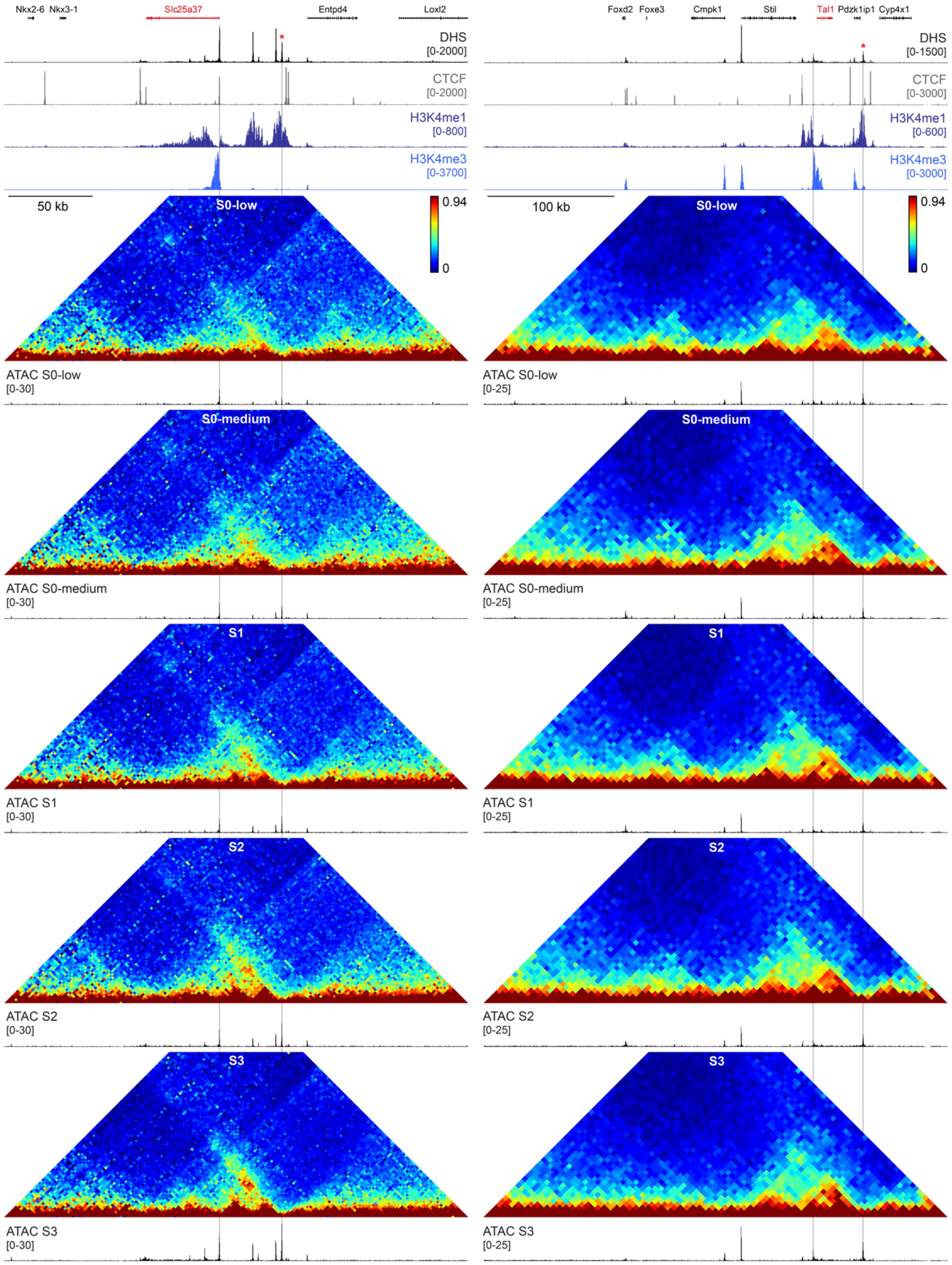

**Supplementary Figure 10: Chromatin interactions and accessibility in the *Slc25a37* and *Tal1* loci through erythroid differentiation. [continued]**

Tiled-C contact matrices spanning the mouse *Slc25a37* (left) and *Tal1* (right) loci in sequential stages of *in vivo* erythroid differentiation at 2 and 5 kb resolution, respectively. Contact frequencies represent normalized, unique interactions from 2 replicates. Matched open chromatin (ATAC) profiles are shown underneath each matrix and represent normalized data from 3 S0-low, S0-medium and S1 replicates and 2 S2 and S3 replicates. Gene annotation (*Slc25a37* and *Tal1* highlighted in red), open chromatin (ATAC) and CTCF, H3K4me3 and H3K4me1 occupancy in mature mouse erythroblasts isolated from spleen are shown at the top. The putative erythroid enhancer elements (defined by increased chromatin accessibility through erythroid differentiation and H3K4me1 occupancy in mature erythroblasts) with which interactions are quantified in Figure 5 are indicated with a red asterisk. Coordinates: chr14:69,776,000-70,050,000 (left) and chr4:114,470,000-114,835,000 (right).
*Slc25a37* encodes for Mitoferrin-1, a mitochondrial iron transporter which is highly expressed in erythroid cells. *Tal1* encodes for T-cell acute lymphoblastic leukemia protein 1, which is an essential transcription factor in hematopoiesis.

**Supplementary Figure 11: Chromatin interactions and accessibility in the *Cd47* and *Cpeb4* loci through erythroid differentiation.**

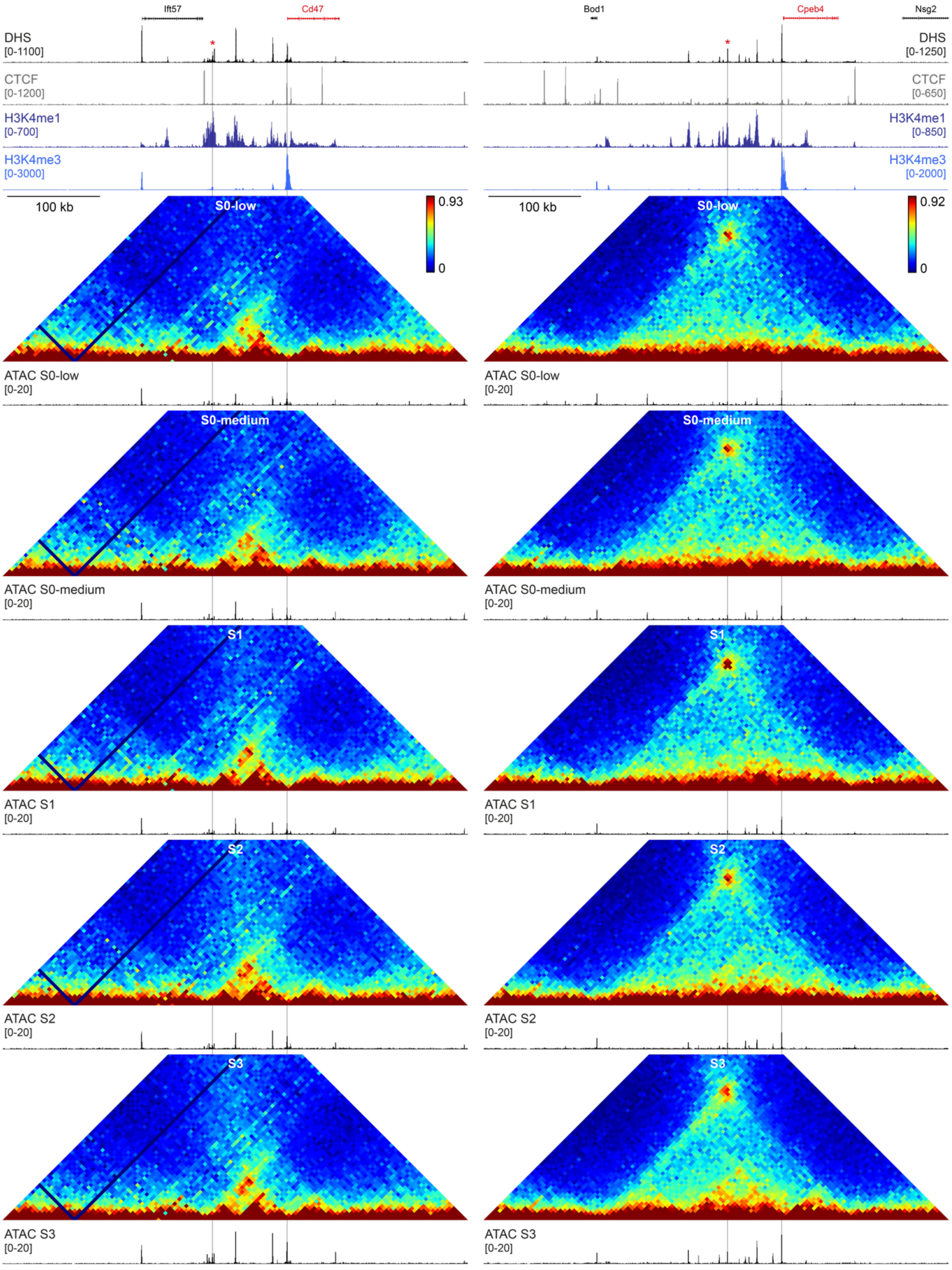

**Supplementary Figure 11: Chromatin interactions and accessibility in the *Cd47* and *Cpeb4* loci through erythroid differentiation. [continued]**

Tiled-C contact matrices spanning the mouse *Cd47* (left) and *Cpeb4* (right) loci in sequential stages of *in vivo* erythroid differentiation at 5 kb resolution. Contact frequencies represent normalized, unique interactions from 2 replicates. Matched open chromatin (ATAC) profiles are shown underneath each matrix and represent normalized data from 3 S0-low, S0-medium and S1 replicates and 2 S2 and S3 replicates. Gene annotation (*Cd47* and *Cpeb4* highlighted in red), open chromatin (ATAC) and CTCF, H3K4me3 and H3K4me1 occupancy in mature mouse erythroblasts isolated from spleen are shown at the top. The putative erythroid enhancer elements (defined by increased chromatin accessibility through erythroid differentiation and H3K4me1 occupancy in mature erythroblasts) with which interactions are quantified in Figure 5 are indicated with a red asterisk. Coordinates: chr16:49,550,000-50,050,000 (left) and chr11:31,450,000-31,950,000 (right).
Cd47 is a transmembrane protein expressed in hematopoietic stem cells and upregulated in erythroid cells to reduce macrophage engulfment. Cpeb4 is an RNA-binding protein which has as essential function in translational control during terminal erythroid development.

**Supplementary Figure 12: Chromatin interactions and accessibility in the *Btg2* locus through erythroid differentiation.**

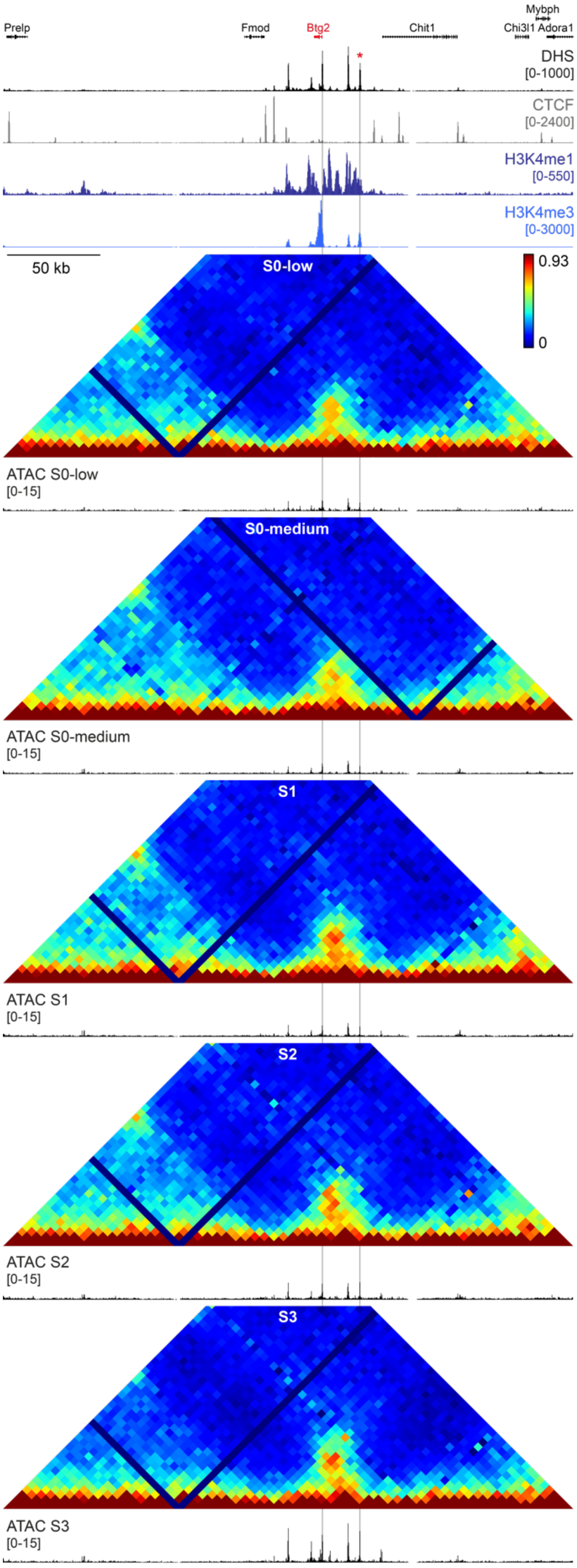

**Supplementary Figure 12: Chromatin interactions and accessibility in the *Btg2* locus through erythroid differentiation. [continued]**

Tiled-C contact matrices spanning the mouse *Btg2* locus in sequential stages of *in vivo* erythroid differentiation at 5 kb resolution. Contact frequencies represent normalized, unique interactions from 2 replicates. Matched open chromatin (ATAC) profiles are shown underneath each matrix and represent normalized data from 3 S0-low, S0-medium and S1 replicates and 2 S2 and S3 replicates. Gene annotation (*Btg2* highlighted in red), open chromatin (ATAC) and CTCF, H3K4me3 and H3K4me1 occupancy in mature mouse erythroblasts isolated from spleen are shown at the top. The putative erythroid enhancer elements (defined by increased chromatin accessibility through erythroid differentiation and H3K4me1 occupancy in mature erythroblasts) with which interactions are quantified in Figure 5 are indicated with a red asterisk. Coordinates: chr1:135,805,000-136,110,000.
*Btg2* (B-cell translocation gene 2) is a tumor suppressor gene and has an important role in hematopoietic differentiation.

**Supplementary Table 1: Comparison of Hi-C and Tiled-C data.**

| | Read pairs | Contacts ($\alpha$-globin region) | Contacts per restriction fragment |
|---|---|---|---|
| **Hi-C ES** | 7,260,480,082 | 3,518,538 | 449 |
| **Tiled-C ES** | 379,033,076 | 97,807,262 | 12,490 |
| **Tiled-C Erythroid** | 293,410,431 | 60,515,876 | 7,728 |

Overview of the number of sequencing reads and detected contacts in the $\alpha$-globin region (chr11:29,902,000-33,228,000) in Hi-C data in mouse ES cells (Bonev et al. Cell 2017) and Tiled-C data in mouse ES and primary erythroid cells. Numbers are derived from pooled data of 4 and 3 replicates for the Hi-C and Tiled-C data, respectively, and contacts represent filtered, non-duplicated contact counts.

**Supplementary Table 2: Comparison of many vs many 3C techniques.**

| | Cell number | Library | Restriction enzyme | Enrichment | Region | Sequencing depth | Resolution |
|---|---|---|---|---|---|---|---|
| **5C** | 100M | 3C library | EcoRI | PCR primers | 500 kb | 550K* (raw) | ** |
| **T2C** | 10M | 3C library | HindIII/ BglII | Microarray; restriction site centered | 2.1 Mb | 52M–85M (raw) | ** |
| **cHi-C** | 20M | Hi-C library | HindIII | 120 bp RNA; not restriction site centered | 3.3 Mb | 62M–99M (mapped) | 10 kb |
| **Hi-C$^2$** | 2–5M | Hi-C library | MboI | 120 bp RNA; restriction site centered | 6 Mb | 600K–60M (raw) | 5–10 kb |
| **Tiled-C** | 2K–10M | 3C library | DpnII | 70 bp DNA; restriction site centered | 1 Mb | 19M-31M (raw) | 2 kb |

Comparison of current many vs many 3C techniques, which generate contact matrices for regions of interest, including Chromosome Conformation Capture Carbon Copy (5C; Dostie et al. Genome Research 2006), Targeted Chromatin Capture (T2C; Kolovos et al. Epigenetics & Chromatin 2014), Capture Hi-C (cHi-C; Dryden et al. Genome Research 2014), HYbrid Capture Hi-C (Hi-C$^2$; Sanborn et al. PNAS 2015) and Tiled-C.

* The original 5C protocol used microarrays or single-molecule pyrosequencing using 454-technology. It is possible to combine 5C with next-generation sequencing at larger sequencing depth to achieve higher resolution.

** 5C and T2C data can be displayed at resolution of individual restriction fragments, which is 2–6 kb for restriction enzymes recognizing a 6 bp sequence. However, the data are then not represented in Hi-C-like contact matrices with equal bin sizes. When 5C data are represented in contact matrices, the data are usually binned in 20–50 kb bins.

**Supplementary Table 3: Comparison of Tiled-C data across low-input primary erythroid samples.**

| Cells | Read pairs | Contacts ($\alpha$-globin region) | Contacts per restriction fragment |
|---|---|---|---|
| **100,000** | 58,362,991 | 6,108,837 | 780 |
| **20,000** | 57,242,048 | 5,782,472 | 738 |
| **10,000** | 55,480,867 | 3,251,628 | 415 |
| **5,000** | 56,476,873 | 1,769,051 | 226 |
| **2,000** | 33,898,548 | 926,504 | 118 |

Overview of the number of sequencing reads and detected contacts in the $\alpha$-globin region (chr11:29,902,000-33,228,000) in Tiled-C data generated from low-input aliquots of primary erythroid cells. Numbers are derived from pooled data of 3 replicates and contacts represent filtered, non-duplicated contact counts.

**Supplementary Table 4: Overview of Tiled-C experiments and capture oligonucleotides.**

| Region | Coordinates | Size (bp) | Oligo number | Cell types |
|---|---|---|---|---|
| α-globin | chr11:29,902,950-33,440,011 | 3,537,061 | 8,571 | Erythroid, ES, Erythroid Prog. |
| Pnpo | chr11:96,470,318-97,399,355 | 929,037 | 2,480 | Erythroid, ES |
| Sox2 | chr3:33,200,154-36,559,869 | 3,359,715 | 8,266 | Erythroid, ES |
| Tal1 | chr4:113,693,194-116,236,558 | 2,543,364 | 6,200 | Erythroid, ES |
| Adra2b | chr2:126,937,437-127,444,583 | 507,146 | 1,519 | Erythroid Prog. |
| Btg2 | chr1:135,800,202-136,109,764 | 309,562 | 909 | Erythroid Prog. |
| Cd47 | chr16:49,545,434-50,053,629 | 508,195 | 1,525 | Erythroid Prog. |
| Cd59b | chr2:103,690,279-103,998,406 | 308,127 | 845 | Erythroid Prog. |
| Epb42 | chr2:120,646,951-121,154,825 | 507,874 | 1,338 | Erythroid Prog. |
| GM867 | chr10:75,088,422-75,594,935 | 506,513 | 1,354 | Erythroid Prog. |
| Hemgn | chr4:46,295,673-46,604,534 | 308,861 | 837 | Erythroid Prog. |
| Mgll | chr6:88,615,170-88,924,985 | 309,815 | 1,014 | Erythroid Prog. |
| Mxd3 | chr13:55,255,664-55,764,675 | 509,011 | 1,353 | Erythroid Prog. |
| Plpp1 | chr13:113,405,249-113,814,811 | 409,562 | 1,077 | Erythroid Prog. |
| Sh3tc2 | chr18:61,926,721-62,433,360 | 506,639 | 1,581 | Erythroid Prog. |
| Slc25a37 | chr14:69,745,662-70,054,416 | 308,754 | 974 | Erythroid Prog. |
| Slc4a1 | chr11:101,976,404-102,483,902 | 507,498 | 1,388 | Erythroid Prog. |
| Sphk1 | chr11:116,265,335-116,574,646 | 309,311 | 835 | Erythroid Prog. |
| Tal1 | chr4:114,465,794-114,974,432 | 508,638 | 1,221 | Erythroid Prog. |
| Tinagl1 | chr4:129,755,887-130,064,932 | 309,045 | 917 | Erythroid Prog. |

Overview of regions targeted by Tiled-C experiments in mature erythroid and ES cells (top) and erythroid progenitor populations (bottom). We have focused our analyses on the regions containing interactions between (putative) enhancers and promoters and included only these regions in the manuscript. The complete datasets are available on GEO.

**Supplementary Table 5: Comparison of Tiled-C data generated using single- and double-stranded oligonucleotides.**

| Oligo | Cell number | Read pairs | Contacts (α-globin region) | Contacts per restriction fragment |
|---|---|---|---|---|
| **Single-stranded** | 100,000 | 58,362,991 | 6,108,837 | 780 |
| **Double-stranded** | 100,000 | 58,636,489 | 5,400,851 | 690 |
| **Single-stranded** | 10,000 | 55,480,867 | 3,251,628 | 415 |
| **Double-stranded** | 10,000 | 56,129,518 | 3,190,934 | 407 |

Overview of the number of sequencing reads and detected contacts in the α-globin region (chr11:29,902,000-33,228,000) in Tiled-C data generated in erythroid cells using single- and double-stranded oligonucleotides. Numbers are derived from pooled data of 3 replicates and contacts represent filtered, non-duplicated contact counts.
